# DNA damage response signatures are associated with frontline chemotherapy response and routes of tumor evolution in extensive stage small cell lung cancer

**DOI:** 10.1101/2024.07.29.605595

**Authors:** Benjamin B. Morris, Simon Heeke, Yuanxin Xi, Lixia Diao, Qi Wang, Pedro Rocha, Edurne Arriola, Myung Chang Lee, Darren R. Tyson, Kyle Concannon, Kavya Ramkumar, C. Allison Stewart, Robert J. Cardnell, Runsheng Wang, Vito Quaranta, Jing Wang, John V. Heymach, Barzin Y. Nabet, David S. Shames, Carl M. Gay, Lauren A. Byers

**Affiliations:** Department of Thoracic/Head and Neck Medical Oncology, The University of Texas M.D. Anderson Cancer Center, Houston, Texas, USA; Department of Bioinformatics and Computational Biology, The University of Texas M.D. Anderson Cancer Center, Houston, Texas, USA; Medical Oncology Department, Vall d’Hebron University Hospital, Barcelona, Spain; Medical Oncology Department, Hospital del Mar, Barcelona, Spain; Department of Oncology Biomarker Development, Genentech Inc., South San Francisco, California, USA; Department of Pharmacology, Vanderbilt University School of Medicine, Nashville, Tennessee, USA; Department of Hematology/Oncology, The University of Texas M.D. Anderson Cancer Center, Houston, Texas, USA; Department of Biochemistry, Vanderbilt University School of Medicine, Nashville, Tennessee, USA

## Abstract

**Introduction:** A hallmark of small cell lung cancer (SCLC) is its recalcitrance to therapy. While most SCLCs respond to frontline therapy, resistance inevitably develops. Identifying phenotypes potentiating chemoresistance and immune evasion is a crucial unmet need. Previous reports have linked upregulation of the DNA damage response (DDR) machinery to chemoresistance and immune evasion across cancers. However, it is unknown if SCLCs exhibit distinct DDR phenotypes.

**Methods:** To study SCLC DDR phenotypes, we developed a new DDR gene analysis method and applied it to SCLC clinical samples, *in vitro*, and *in vivo* model systems. We then investigated how DDR regulation is associated with SCLC biology, chemotherapy response, and tumor evolution following therapy.

**Results:** Using multi-omic profiling, we demonstrate that SCLC tumors cluster into three DDR phenotypes with unique molecular features. Hallmarks of these DDR clusters include differential expression of DNA repair genes, increased replication stress, and heightened G2/M cell cycle arrest. SCLCs with elevated DDR phenotypes exhibit increased neuroendocrine features and decreased “inflamed” biomarkers, both within and across SCLC subtypes. Treatment naive DDR status identified SCLC patients with different responses to frontline chemotherapy. Tumors with initial DDR Intermediate and DDR High phenotypes demonstrated greater tendency for subtype switching and emergence of heterogeneous phenotypes following treatment.

**Conclusions:** We establish that SCLC can be classified into one of three distinct, clinically relevant DDR clusters. Our data demonstrates that DDR status plays a key role in shaping SCLC phenotypes, chemotherapy response, and patterns of tumor evolution. Future work targeting DDR specific phenotypes will be instrumental in improving patient outcomes.

## Introduction

Small cell lung cancer (SCLC) is a high-grade neuroendocrine malignancy and the most lethal form of lung cancer [1]. This poor prognosis is due in large part to SCLC’s recalcitrance to therapy. Unlike most non-small cell lung cancers (NSCLCs), SCLCs lack targetable oncogenic driver alterations and instead develop through dual inactivation of *TP53* and *RB1* tumor suppressor genes [2, 3]. In lieu of oncogene-directed targeted therapies, a chemotherapy doublet of etoposide and platinum-based chemotherapy (EP) is the backbone of SCLC frontline therapy. While most SCLCs are initially exquisitely sensitive to chemotherapy, most tumors recur and rapidly progress as chemoresistant disease soon after initial treatment [1]. The addition of anti-PDL1 immunotherapy to EP chemotherapy has improved outcomes for extensive stage (ES) SCLC patients [4]. However, frontline chemoimmunotherapy does not produce durable disease control for the vast majority of ES-SCLC patients. Identifying and characterizing biologic programs potentiating chemoresistance and immune evasion is a crucial unmet need in SCLC.

Research from our group and others has demonstrated that transcriptional profiling can identify clinically relevant, molecularly distinct subtypes of SCLC [5–7]. Using unsupervised clustering and large clinical datasets, Gay et al. found that SCLC is composed of four distinct subtypes marked by expression of lineage-specific transcription factors or characterized by “inflamed” biomarkers [5]. Clinically, “inflamed” tumors, which account for roughly 20% of *de novo* SCLC, tend to derive more durable benefit from frontline chemoimmunotherapy. Despite this knowledge, it is unknown why the overwhelming majority of ES-SCLCs do not respond durably following frontline treatment.

Studies have shown that increased expression of DNA repair genes is associated with chemotherapy resistance and immune cold tumor microenvironments across cancers [8–11]. In SCLC, recent studies have identified an inverse correlation between DNA repair gene expression and “inflamed” biomarkers in relapsed tumors [12]. However, it is unknown if SCLCs exhibit distinct DNA damage response (DDR) phenotypes and if these phenotypes identify tumors with unique molecular features not currently captured by SCLC subtypes alone.

We hypothesized that differential regulation of the DDR machinery would identify biologically distinct, clinically relevant clusters of SCLC tumors. To test our hypothesis, we developed a new DDR gene expression analysis method and applied it to SCLC clinical samples, *in vitro*, and *in vivo* SCLC model systems. Using this method, we investigated how expression of the DDR machinery is associated with neuroendocrine biology, immune evasion, frontline chemotherapy response, and tumor evolution in SCLC.

## Materials and Methods

### Molecular profiling data

We analyzed bulk RNA sequencing data for SCLC tumor samples from the MD Anderson GEMINI cohort [13], the IMpower133 clinical trial [4, 7], SCLC cell lines [14], and SCLC and high grade neuroendocrine (hgNEC) patient derived xenografts (PDX) and circulating tumor cell derived xenografts (CDX) models. Single cell RNA sequencing (scRNAseq) from these PDX/CDX models was also analyzed. SCLC cell line reverse phase proteomic array (RPPA) protein expression data was previously generated by our group and the Broad Institute [14, 15]. GEMINI, IMpower133, and SCLC PDX/CDX RNA sequencing data are presented as normalized transcripts per million (TPM) values. SCLC cell line gene expression data are presented as z-scores, as deposited by Tlemsani et al. [14]. RPPA data are presented as normalized, log2 transformed values. Reduced representation bisulfite sequencing (RRBS) methylation data analyzed in this study were previously generated by Heeke et al. [13].

### Weighted Expression Score (WE Score) Analysis Method

WE scores were calculated to evaluate transcriptional regulation of ten distinct DDR pathways in SCLC samples (**Supplementary Figure 1A**). Gene expression z-scores were calculated for 130 DDR genes functioning across the 10 pathways. Z-score values were used to prevent differences in transcript abundance across genes from skewing pathway regulation scores. WE scores were generated by multiplying gene expression z score values by gene-specific essentiality scaling factor (ESF) values to account for effector importance to pathway function (**Supplementary Figure 1B**). ESF values were assigned following extensive literature review and are based on DDR gene essentiality in genetic knockout studies and familial cancer syndromes [16–25]. DDR gene ESF values are listed in **Supplementary Table 1**. These products were summed and divided by the total number of genes in each pathway to generate raw WE scores. Raw WE scores were then scaled across samples. The fviz_nbclust function in the factoextra R package was used to determine the optimal number of *k* clusters by minimizing the total within sum of squares variance between clusters. A maximum of k = 10 clusters were considered. Samples were clustered using the ComplexHeatmap R package and the statistically defined optimal number of *k* clusters.

### SCLC Subtype Assignment

IMpower133 tumor samples and SCLC cell lines were subtyped using our previously described non-negative matrix factorization (NMF) approach and 1300 gene signature [5]. GEMINI tumor samples and SCLC PDX/CDX models were subtyped using our new SCLC Gene-Ratio Classifier (SCLC-GRC) [13]. As our SCLC-GRC method was trained using IMpower133 samples, this approach could not be used to subtype IMpower133 samples.

### Differential gene expression and quantitative set analysis for gene expression (QuSAGE)

Differential gene expression and quantitative set analysis for gene expression (QuSAGE) was used to identify transcriptional hallmarks of IMpower133 DDR clusters. Genes differentially expressed between DDR clusters were identified using pairwise comparisons and the limma R package [7, 26]. Gene set analyses were performed using the QuSAGE R package [7, 27].

### Immune and stromal cell infiltration estimation

The tidyestimate R package was used to estimate bulk immune and stromal cell infiltration in GEMINI and IMpower133 samples [28].

### RepStress and Neuroendocrine Score (NE Score) Analyses

RepStress and Neuroendocrine Scores were calculated as described previously [11, 29].

### scRNAseq Analysis

SCLC PDX/CDX tumors were processed and sequenced using the 10X genomics platform as described previously [30]. The Cell Ranger v3.0.1 pipeline was used to process raw reads to UMI counts [31]. Samples from different sequencing batches were integrated and normalized using Seurat’s SCT integration pipeline [32]. Cells expressing less than 200 genes or those with mitochondrial reads accounting for >20% of total reads were removed. Principal component analysis (PCA) and UMAP transformations were used for dimensionality reduction with Seurat. Cell cycle states were determined using Seurat’s “CellCycleScoring” function. Binary expression plots were generated using in-house R scripts.

### RNA Velocity Analysis

scRNAseq data from PDX model SC53 were assessed for stable and transient states based on RNA velocity using scvelo and cellrank Python packages [33–35]. Cells were filtered for quality as described previously [5]. The top 2000 highly variable genes were selected and used for downstream analyses. PCA was applied and the first 30 components were used to identify the 30 nearest neighbors of each cell (kNN). The latent gene space was subjected to t-distributed stochastic network embedding (t-SNE) and clusters were identified using the Leiden algorithm. A model of transcriptional dynamics of splicing kinetics is solved in a likelihood-based expectation maximization by estimating transcriptional state and cell-internal latent time to learn the unspliced/spliced phase trajectory for each gene. This was performed using scvelo’s recover_dynamics in stochastic mode. Driver genes for transitions across states (Leiden clusters) were detected by their high likelihoods in the dynamic model. Random walk simulations were performed to mimic temporal transitions and likely resultant end states using 100 cells randomly placed within each Leiden cluster and proceeding until the system reached steady state. The WEB-based Gene SeT AnaLysis Toolkit (WebGestalt) platform was used to conduct over-representation analyses in order to identify biological processes significantly upregulated in single cell cluster 8 [36].

### Subtype Switching Analysis

Patient matched longitudinal liquid biopsy samples were subtyped using circulating tumor DNA methylation profiles and our new SCLC DNA methylation classifier (SCLC-DMC) [13]. Patients whose disease progression liquid biopsy was classified as a different SCLC subtype compared to their matched treatment naïve liquid biopsy were deemed to have ‘subtype switched’ following frontline chemoimmunotherapy.

### Statistical Analyses

Statistical and bioinformatic analyses were performed in R version 4.2. Wilcoxon tests were used to compare gene expression and protein expression data across DDR clusters. All *p*-values are derived from Wilcoxon tests unless otherwise specified. Boxplots visualize data median as well as 1^st^ and 3^rd^ interquartile ranges. Median overall survival (OS) and progression free survival (PFS) were calculated using the Kaplan-Meier method. The IMpower133 trial was not powered to detect statistically significant differences in molecular subtypes within trial arms. Chi-square tests were used to test for significant enrichment patterns when comparing SCLC subtype and DDR cluster assignments. Chi-square tests were also performed with simulation (2000 replicates) to account for small *n* across some assignment categories. Paired Wilcoxon tests were used to compare promoter methylation values for our patient-matched baseline and progression liquid biopsy cohort. *p*-values <0.05 were considered statistically significant.

## Results

### SCLC clusters into three distinct DDR clusters with unique molecular features

To study SCLC DDR phenotypes, we applied a new DDR analysis method to two SCLC clinical datasets, MD Anderson GEMINI (n = 85) and IMpower133 (n = 271) (**Methods, Supplementary Figure 1**). GEMINI SCLC samples were largely treatment naïve, ES-SCLC tumor samples collected at MD Anderson Cancer Center [13]. IMpower133 samples were exclusively treatment naïve, ES-SCLC tumor samples collected as part of the IMpower133 Phase III clinical trial [4, 7]. Using unsupervised k-means clustering, we identified three statistically defined clusters of GEMINI SCLC tumors demarcated by DDR phenotypes (**Figure 1A**). DDR Low tumors were characterized by lowest expression of almost all DDR pathways. DDR Intermediate tumors exhibited a heterogeneous pattern of intermediate to high expression of various DDR pathways. DDR High tumors displayed strong upregulation of almost all DDR pathways. To validate these findings, we applied our DDR analysis method to an independent dataset of IMpower133 tumor samples. As in our GEMINI dataset, unsupervised k-means clustering again identified three statistically defined clusters of IMpower133 tumors as defined by unique DDR phenotypes (**Figure 1B**). Across both datasets, we found a highly similar distribution of samples in each DDR cluster (**Supplementary Figure 2A-B**). Inspection of our DDR clusters confirmed that our method indeed identified groups of SCLC tumors with striking differences in individual DDR gene expression patterns (**Supplementary Figure 2C-D**). It should be noted that our method clusters tumors with similar global DDR phenotypes by simultaneously evaluating expression of 130 genes functioning across 10 distinct DNA repair pathways. Thus, our method represents a significant divergence from single gene stratification approaches where expression of a single gene differentiates samples.

**Figure 1:**
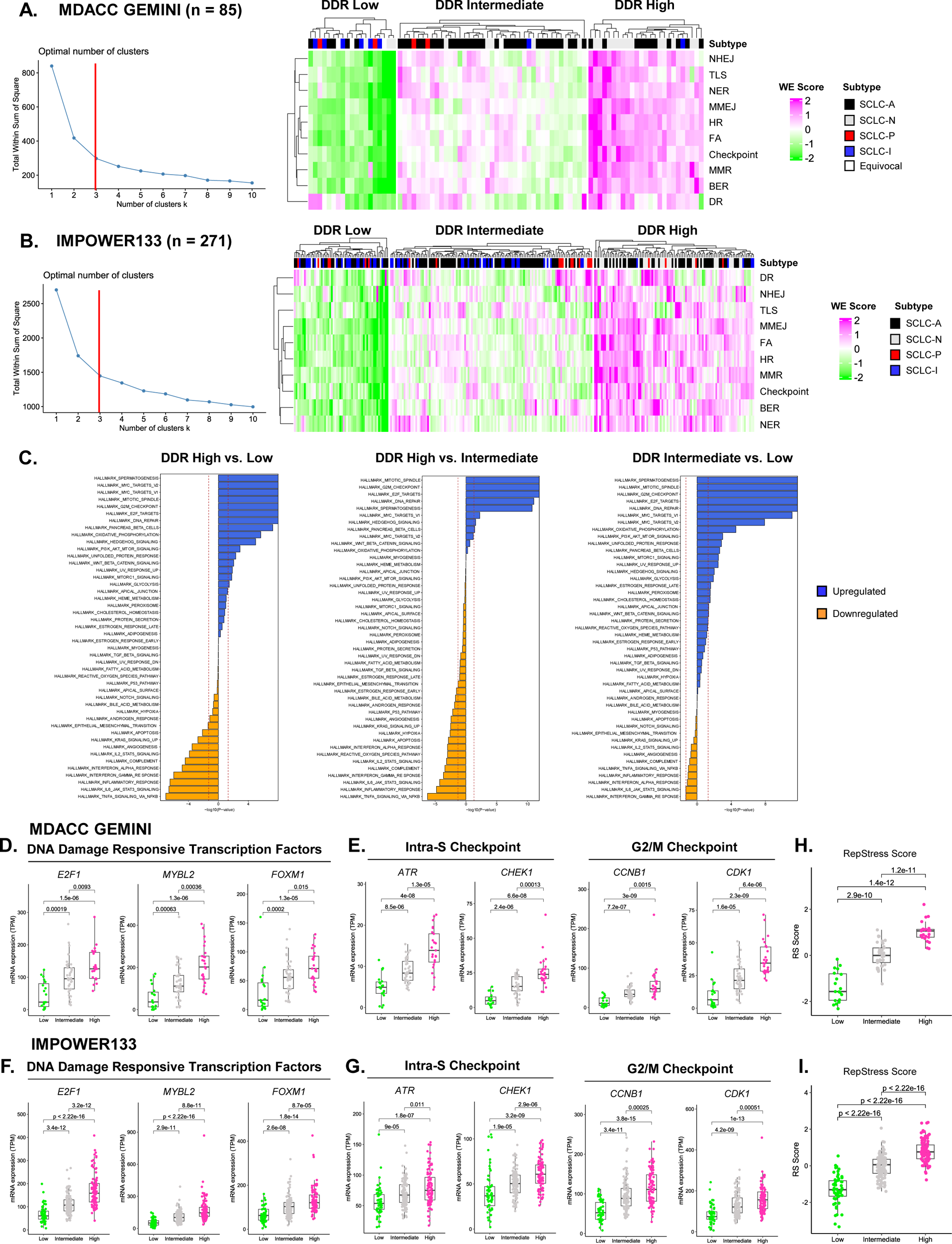
SCLC clusters into three distinct DDR clusters with unique molecular features. **A.** MD Anderson GEMINI optimal number of k clusters elbow plot and DDR cluster heatmap. **B.** Optimal number of k clusters elbow plot and DDR cluster heatmap for the IMPOWER133 cohort. **C.** IMpower133 DDR cluster differential gene expression and quantitative set analysis (QuSAGE) hallmark pathway results. **D.** GEMINI DDR cluster DNA damage responsive transcription factor expression. **E.** GEMINI DDR cluster intra-S and G2/M cell cycle checkpoint machinery expression. **F.** Expression of DNA damage responsive transcription factors in IMPOWER133 DDR clusters. **G.** Expression of intra-S and G2/M cell cycle checkpoint machinery in IMPOWER133 DDR clusters. **H-I.** GEMINI and IMPOWER133 DDR cluster RepStress scores. **A-B.** NHEJ: Non-homologous end-joining. TLS: Translesion synthesis. NER: Nucleotide excision repair. MMEJ: Microhomology mediated end-joining. HR: Homologous recombination. FA: Fanconi Anemia. Checkpoint: Damage sensing and signaling. MMR: Mismatch repair. BER: Base excision repair. DR: Direct reversal repair.

We next sought to identify unique molecular features of tumors with differing DDR status. Using QuSAGE analysis, we found that DDR High tumors highly express hallmark pathways controlling DNA repair, E2F transcriptional targets, MYC transcriptional targets, and the G2/M cell cycle checkpoint, compared to DDR Low (**Figure 1C**). DDR Low tumors, meanwhile, were characterized by upregulation of pathways controlling apoptosis, interferon alpha signaling, interferon gamma signaling, epithelial to mesenchymal transition (EMT), TNF signaling, immune cell trafficking, and immune cell function, compared to DDR High (**Figure 1C, Supplementary Figure 3**). DDR Intermediate tumors showed evidence of hybrid phenotypes marked by upregulation of DNA repair, cell cycle checkpoint, and inflammatory programs when compared to either DDR High or DDR Low tumors. Collectively, these data indicate that SCLC tumors with different DDR status exhibit a spectrum of DNA repair and inflammatory phenotypes.

Given that gene set enrichment analysis identified E2F transcriptional targets and cell cycle checkpoint pathways as key pathways differentially expressed across DDR clusters, we next investigated expression of individual effectors controlling these processes. Across both GEMINI and IMpower133 datasets, we confirmed that expression of DNA damage responsive transcription factors—including E2F1—significantly differed across DDR clusters (**Figure 1D, 1F**). It should be noted that these genes were not included in our DDR analysis method and represent orthogonal validation of unique DDR cluster biology. In addition, we also confirmed that expression of intra-S and G2/M cell cycle checkpoint machinery effectors significantly increased in a DDR specific manner (**Figure 1E, 1G**). As DDR clusters were marked by differential expression of DNA repair and cell cycle checkpoint genes, we hypothesized that SCLC tumors across these clusters experience varying levels of replication stress. Previous studies have demonstrated that the RepStress transcriptional signature reliably identified differences in cellular replication stress and endogenous DNA damage [11]. To test our hypothesis, we calculated RepStress scores for SCLC tumor samples in GEMINI and IMpower133 datasets. As expected, we found that a hallmark of different DDR status is higher levels of replication stress (**Figure 1H, 1I)**. Collectively, these data demonstrate that SCLC is composed of three biologically distinct DDR clusters with unique molecular features.

### SCLC DDR clusters are recapitulated in both in vitro and in vivo model systems

To extend our findings, we applied our DDR analysis method to a cohort of SCLC cell line models (n = 116) [14]. As in human SCLC samples, unsupervised k-means clustering identified three statistically defined clusters of SCLC cell lines as described by DDR phenotypes (**Figure 2A, Supplementary Figure 4**). We observed a similar distribution of cell line models across DDR clusters, with a slight enrichment of DDR High models compared to that seen in human samples (**Supplementary Figure 5A**). Consistent with our previous findings, cell line DDR clusters exhibited differences in DNA damage responsive transcription factor and cell cycle checkpoint machinery transcripts (**Supplementary Figure 5B**). Additionally, we observed a dramatic difference in replication stress across cell line DDR clusters (**Figure 2B**). Beyond different transcriptional phenotypes, we found that SCLC cell line DDR clusters were marked by significant differences in protein expression. Protein expression of DNA damage responsive transcription factors, DNA damage sensing kinases, and replication stress mediators were significantly different across DDR clusters (**Figure 2C-D**). Additionally, expression of DNA repair proteins that function in specific repair pathways—MSH2 in MMR and XRCC1 in BER— increased across DDR clusters. Furthermore, we found expression of proteins that indirectly promote DNA repair also increased across DDR clusters (**Figure 2E**). Interestingly, we found no difference in SLFN11 protein expression, which has previously been reported to highly correlate with DNA repair gene expression in SCLC (**Supplementary Figure 5C**). Beyond DNA repair proteins, we found that cell line DDR clusters exhibited differential expression of key cell cycle checkpoint regulators (**Figure 2F, Supplementary Figure 5D**). Notably, we observed a concerted upregulation of multiple proteins controlling G2/M cell cycle arrest following DNA damage in DDR High and Intermediate clusters compared to DDR Low (**Figure 2F**). This differential expression of G2/M checkpoint proteins was independent of *MYC* expression (**Supplementary Figure 5E**).

**Figure 2:**
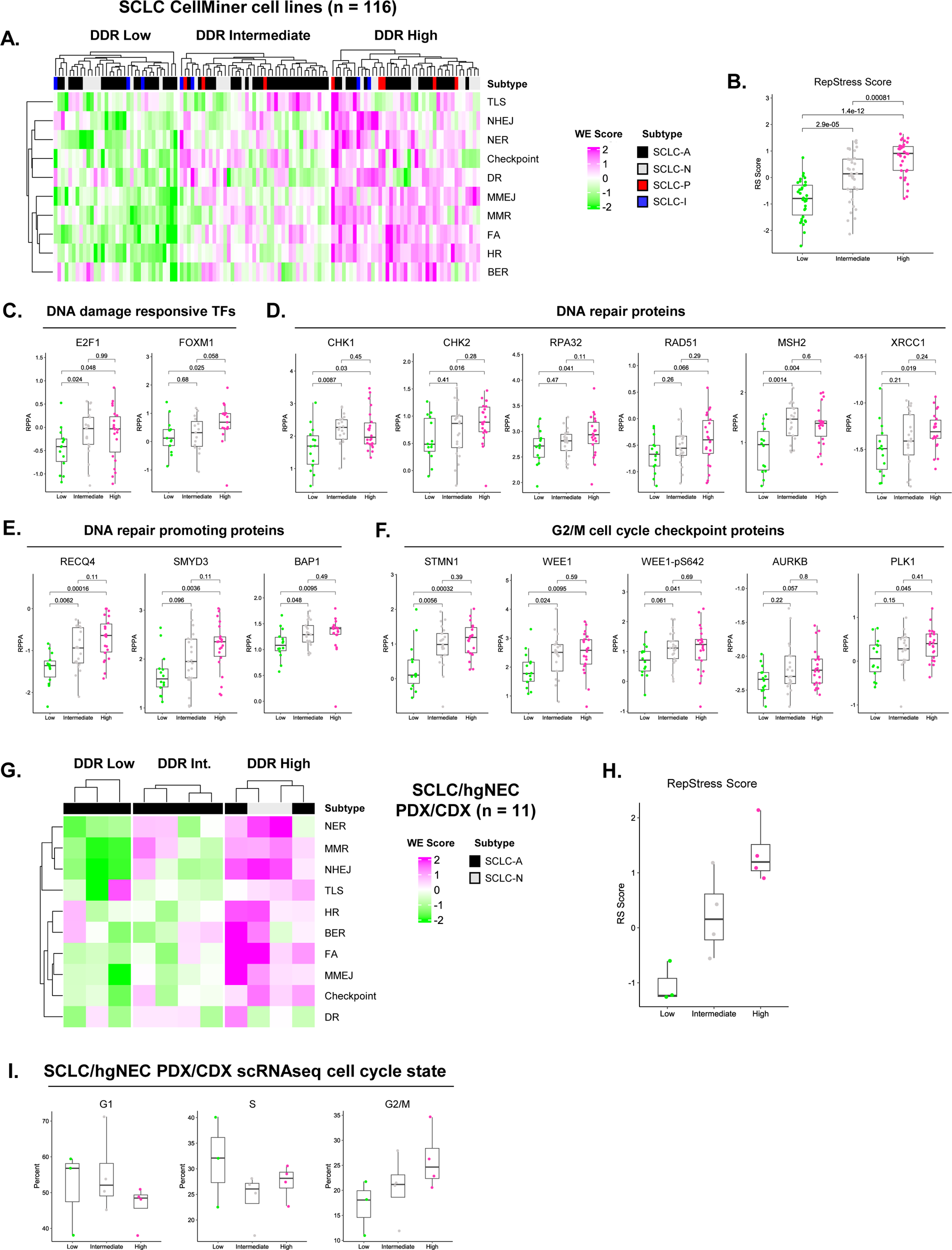
SCLC DDR clusters are recapitulated in both *in vitro* and *in vivo* model systems. **A.** DDR cluster heatmap for SCLC CellMiner cell line models. **B.** SCLC cell line DDR cluster RepStress scores. **C.** SCLC cell line DNA damage responsive transcription factor protein expression. **D.** DNA repair protein expression across SCLC cell line DDR clusters. **E.** Protein expression of DNA repair promoting effectors across DDR clusters. **F.** G2/M cell cycle checkpoint effector protein expression in cell line DDR clusters. **G.** DDR cluster heatmap for SCLC/hgNEC PDX and CDX models. **H.** SCLC/hgNEC PDX/CDX DDR cluster RepStress scores. **I.** SCLC/hgNEC PDX/CDX DDR cluster scRNAseq cell cycle state distribution.

We next applied our DDR analysis method to SCLC PDXs/CDXs. This analysis confirmed that SCLC PDX/CDX models also exhibit three distinct DDR phenotypes (**Figure 2G**). Like human SCLC and cell lines, PDX/CDX DDR clusters were marked by increasing levels of key transcripts and differing levels of replication stress (**Supplementary Figure 6A**, **Figure 2H**). Given that DDR clusters exhibit increased replication stress and markers of G2/M cell cycle arrest, we hypothesized that tumors with different DDR status have different cell cycle state distributions. Using scRNAseq data, we found that increasing DDR status identifies tumors with increasing levels of single tumor cells in G2/M *in vivo* (**Figure 2I, Supplementary Figure 6B**). These data demonstrate that SCLC DDR clusters and their unique molecular features are robustly recapitulated in *in vitro* and *in vivo* model systems.

### SCLC DDR status is significantly associated with SCLC subtypes and neuroendocrine features

Recent studies have shown that SCLC is composed of four subtypes defined by expression of lineage-specific transcription factors or “inflamed” biomarkers [5, 7]. The relationship between these established subtypes and DDR status is unknown. In both GEMINI and IMpower133 datasets, we found DDR status was significantly associated with SCLC subtypes (**Figure 3A-D**). All DDR clusters were composed of tumors from multiple subtypes, with individual subtypes being significantly enriched across different DDR clusters (**Figure 3A, 3C**). Specifically, SCLC-N tumors were enriched in DDR High, SCLC-A in DDR Intermediate, and SCLC-P and SCLC-I in DDR Low.

**Figure 3:**
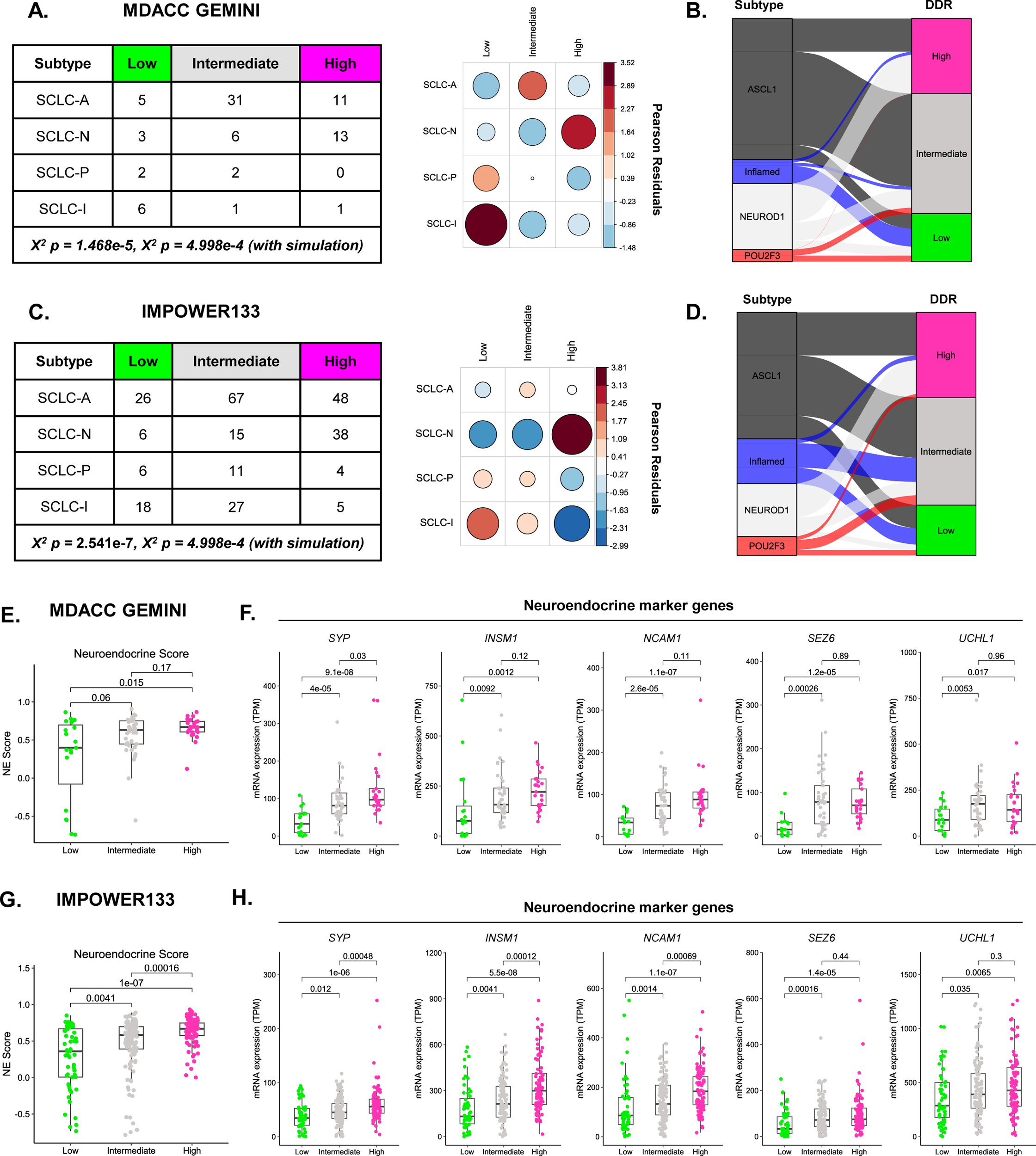
SCLC DDR status is significantly associated with SCLC subtypes and neuroendocrine features. **A.** GEMINI DDR cluster and SCLC subtype assignment table where values listed represent number of patients. *X*^2^ Pearson residual dot plot comparing DDR cluster and SCLC subtype assignments for GEMINI samples. **B.** Alluvial plot of GEMINI DDR cluster composition by SCLC subtype. **C.** IMPOWER133 DDR cluster and SCLC subtype assignment table. Table values represent number of patients. *X*^2^ Pearson residual dot plot comparing DDR cluster and SCLC subtype assignments for IMPOWER133 samples. **D.** Alluvial plot of IMPOWER133 DDR cluster composition by SCLC subtype. **E.** GEMINI DDR cluster neuroendocrine score analysis. **F.** GEMINI DDR cluster neuroendocrine and non-neuroendocrine marker gene expression. **G.** IMPOWER133 DDR cluster neuroendocrine score analysis. **H.** IMPOWER133 DDR cluster neuroendocrine and non-neuroendocrine marker gene expression.

Historically, SCLC tumors have been broadly classified based on the presence or absence of neuroendocrine features [29, 37, 38]. To test if DDR status was associated with differing levels of neuroendocrine features, we analyzed neuroendocrine scores which capture expression of lung-specific neuroendocrine and non-neuroendocrine markers [29]. Across both GEMINI and IMpower133 cohorts, we found that NE scores significantly increased across DDR clusters (**Figure 3E, 3G, Supplementary Figure 7**). This DDR-specific association with neuroendocrine features was further demonstrated by increased expression of individual neuroendocrine marker genes (**Figure 3F, 3H**). We further confirmed these findings in both cell line and PDX/CDX models (**Supplementary Figure 8**). Taken together, these data demonstrate that DDR status is significantly associated with SCLC subtypes and increased neuroendocrine features.

### DDR status identifies a spectrum of “inflamed” features within and across SCLC subtypes

When inspecting DDR signature associations with SCLC subtypes, we were struck by the significant depletion of SCLC-I tumors in DDR High clusters (**Figure 3**). We hypothesized that DDR status also identified tumors with a spectrum of “inflamed” features, both within and across SCLC subtypes. As established by Gay et al., a hallmark of SCLC-I tumors is increased immune cell infiltration [5]. Using the ESTIMATE deconvolution algorithm, we found that immune cell and stromal cell infiltration signals significantly decreased across treatment naïve DDR clusters (**Figure 4A-B**) [28]. These findings agree well with our previous observation that pathways controlling inflammatory responses, antigen presentation, immune cell trafficking, and immune cell function are strongly anti-correlated with elevated DDR signatures (**Figure 1C, Supplementary Figure 3**). In further support of our findings, recent analyses have also demonstrated that inflamed tumors can be found within each SCLC subtype [7]. To test if our observation was independent of the more infiltrated SCLC-I subtype, we restricted our analyses to SCLC-A tumors which are the most prevalent subtype of SCLC and are a more neuroendocrine, “immune cold” subset of SCLC [5, 39]. Strikingly, we found that splitting treatment naïve SCLC-A tumors only by DDR status identified tumors with significantly different levels of immune and stromal cell infiltration signals (**Figure 4C**). Beyond immune cell infiltration, we analyzed other known biomarkers of “inflamed” tumors. Our analysis demonstrated that expression of MHC class I and pro-inflammatory cytokine genes regulating cytotoxic immune cell and antigen presenting cell recruitment decreased across DDR clusters within treatment naïve SCLC-A tumors (**Figure 4D-E**). Additionally, we found that expression of immune checkpoint marker *LAG3* increased with increasing DDR status (**Figure 4E**).

**Figure 4:**
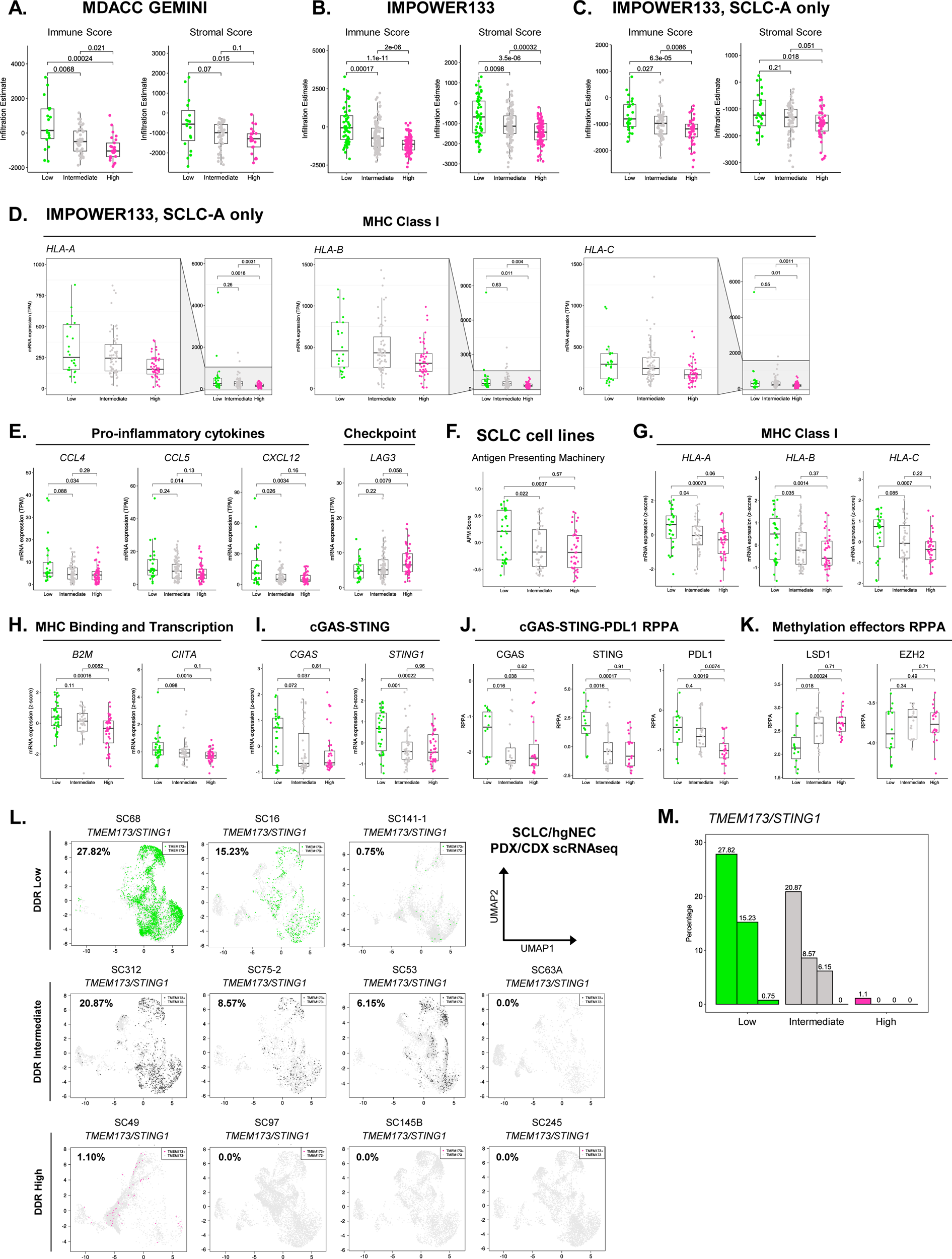
DDR status identifies a spectrum of “inflamed” features both within and across SCLC subtypes. **A.** GEMINI DDR cluster ESTIMATE Immune and Stromal scores. **B.** IMPOWER133 DDR cluster Immune and Stromal scores. **C.** IMPOWER133 SCLC-A only DDR cluster Immune and Stromal scores. **D.** IMPOWER133 SCLC-A only DDR cluster MHC Class I gene expression. **E.** IMPOWER133 SCLC-A DDR cluster pro-inflammatory cytokine and immune suppressive checkpoint effector gene expression. **F.** SCLC cell line DDR cluster Antigen Presenting Machinery scores. **G.** SCLC cell line DDR cluster MHC Class I gene expression. **H.** SCLC cell line DDR cluster B2M and CIITA gene expression. **I.** SCLC cell line DDR cluster cGAS and STING gene expression. **J.** SCLC cell line DDR cluster cGAS, STING1, and PDL1 protein expression. **K.** SCLC cell line DDR cluster LSD1 and EZH2 protein expression. **L.** SCLC/hgNEC PDX/CDX DDR cluster single cell RNAseq *STING1* binary expression plots. **M.** SCLC/hgNEC PDX/CDX DDR scRNAseq *STING1* binary expression summary plot.

To confirm that these observations were independent of tumor purity, we repeated our analyses in SCLC cell lines. This analysis confirmed that pure SCLC tumor cells had significantly decreased antigen presenting machinery expression when stratified by DDR status (**Figure 4F-H**). This decreased MHC Class I expression was correlated with decreased expression of *CIITA*, a transcription factor responsible for driving MHC Class I gene transcription [40]. Studies have shown that expression of the MHC Class I locus can be positively regulated by inflammatory signaling through the cGAS-STING pathway [41]. Given this, we next investigated if cGAS-STING pathway dysregulation was coincident with MHC Class I silencing in DDR clusters. We found that DDR High and Intermediate models had significantly decreased expression of both *cGAS* and *STING1*, compared to DDR Low (**Figure 4I**). This difference was confirmed at the protein level whereas DDR High and Intermediate models had significantly decreased expression of cGAS, STING, and immunomodulatory PDL1 proteins (**Figure 4J**). In addition to cGAS-STING dysregulation, we found that protein expression of the epigenetic regulator LSD1 increased in a DDR specific manner and was correlated with MHC Class I silencing, consistent with recent reports (**Figure 4K**) [6, 42, 43]. Lastly, we analyzed scRNAseq data from SCLC PDX/CDX models to confirm that these differences were also observed in SCLC tumor cells *in vivo*. As expected, we found that expression of *TMEM173/STING1* decreased across DDR clusters in single tumor cells (**Figure 4L-M**). A similar trend was also observed for MHC Class I genes (**Supplementary Figure 9**). Collectively, these data robustly demonstrate that DDR status identifies a spectrum of “inflamed” biology both within and across known SCLC subtypes.

### DDR status identifies tumors with different responses to frontline chemotherapy

Given that SCLC DDR clusters exhibited striking differences in DNA repair and replication stress phenotypes, we hypothesized that DDR status would be associated with different responses to chemotherapy in the clinic. To assess the effect of DDR status on frontline chemotherapy response, we analyzed data from the etoposide + carboplatin arm of the IMPower133 Phase III clinical trial [4]. Interestingly, we found that DDR status associated with different depth of response following frontline chemotherapy, as measured by RECIST criteria [44]. DDR High tumors trended toward strongest depth of initial response, measured by RECIST maximum SLD change, followed by DDR Intermediate, and DDR Low tumors (**Figure 5A**). This difference in initial tumor shrinkage was recapitulated in differences in RECIST best overall response. Rates of response to frontline chemotherapy were increased in tumors with DDR Intermediate and High phenotypes, compared to DDR Low (**Figure 5B, Supplementary Figure 10**). Conversely, DDR Low tumors were enriched for patients whose tumors did not achieve partial or complete radiographic responses. Paradoxically, patients with DDR High and Intermediate disease had numerically shorter OS and PFS, despite experiencing strongest initial tumor reduction following frontline chemotherapy (**Figure 5C, Supplementary Figures 10-11**). This difference was even more pronounced within SCLC-A tumors, where patients with DDR Low tumors had a median OS of 12.65 months (Low vs. Intermediate or High HR = 0.54 (95% CI 0.23-1.28)), compared to 9.85 months for DDR Intermediate, and only 9.49 months in DDR High (**Figure 5D**). These data suggest that DDR status identified SCLC patients with differing responses to frontline chemotherapy.

**Figure 5:**
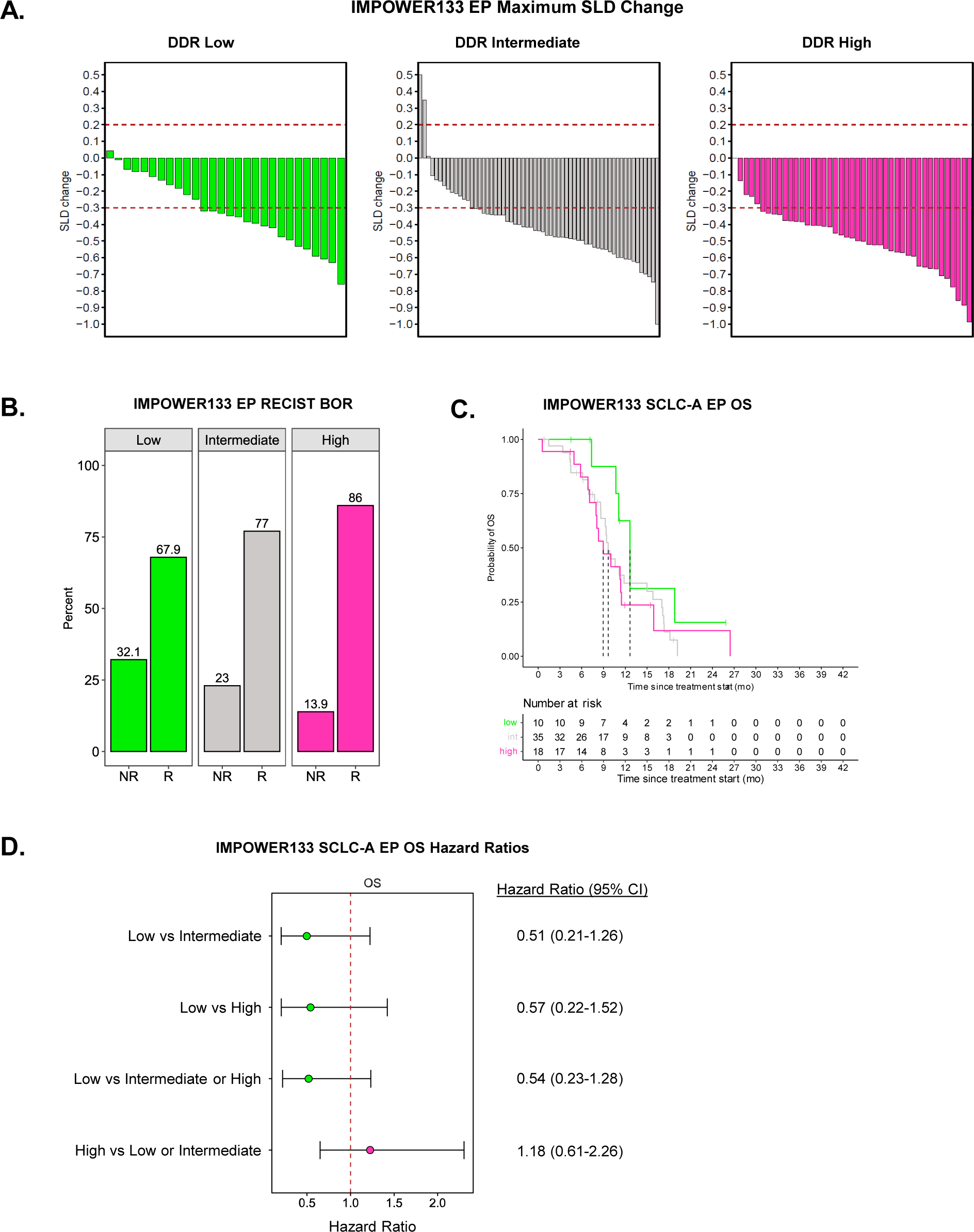
DDR status identifies tumors with different responses to frontline chemotherapy. **A.** Maximum sum of longest dimensions (SLD) change for IMPOWER133 DDR clusters following frontline EP chemotherapy. **B.** RECIST Best Overall Response (BOR) for IMPOWER133 DDR clusters following frontline EP chemotherapy. NR: Non-responder (progressive disease + stable disease). R: Responder (partial response + complete response). **C.** Kaplan Meier plot for IMPOWER133 SCLC-A DDR cluster OS outcomes following frontline EP chemotherapy. **D.** Forrest plot for IMPOWER133 SCLC-A DDR cluster OS outcomes following frontline EP chemotherapy.

### DDR status is linked to subtype switching and unique routes of disease progression following frontline chemoimmunotherapy

Recent studies have demonstrated that SCLCs can ‘switch’ subtypes and evolve to more ‘inflamed’ phenotypes following therapy [13, 30]. Given that SCLC DDR status is linked to distinct tumor microenvironments at baseline, we hypothesized that initial DDR status would influence tumor evolution following frontline chemoimmunotherapy. To test our hypothesis, we analyzed matched treatment naïve tumor tissue, treatment naïve plasma, and plasma collected at disease progression at first recurrence for six ES-SCLC patients (**Figure 6A**). Of these six samples, four tumors underwent subtype switching following frontline therapy—shifting from an SCLC-A state at baseline to an “inflamed” state at progression (**Methods, Figure 6B, Supplementary Figure 12**). Strikingly, patients with inflamed subtype switching tumors did not receive durable benefit from frontline chemoimmunotherapy, with OS and PFS outcomes far inferior to patients with *de novo* SCLC-I tumors (**Supplementary Figure 13**) [5]. All four of these subtype switching tumors were either DDR Intermediate or DDR High. When analyzing these samples further, we found these plastic tumors had some of the lowest initial immune infiltrates across a third independent cohort of treatment naïve SCLC tumors (**Figure 6C**) [13]. These data suggests that DDR-specific immune cell poor microenvironments may be linked to the ability of SCLC cells to progress to an “inflamed” state but not receive durable benefit from frontline chemoimmunotherapy.

**Figure 6:**
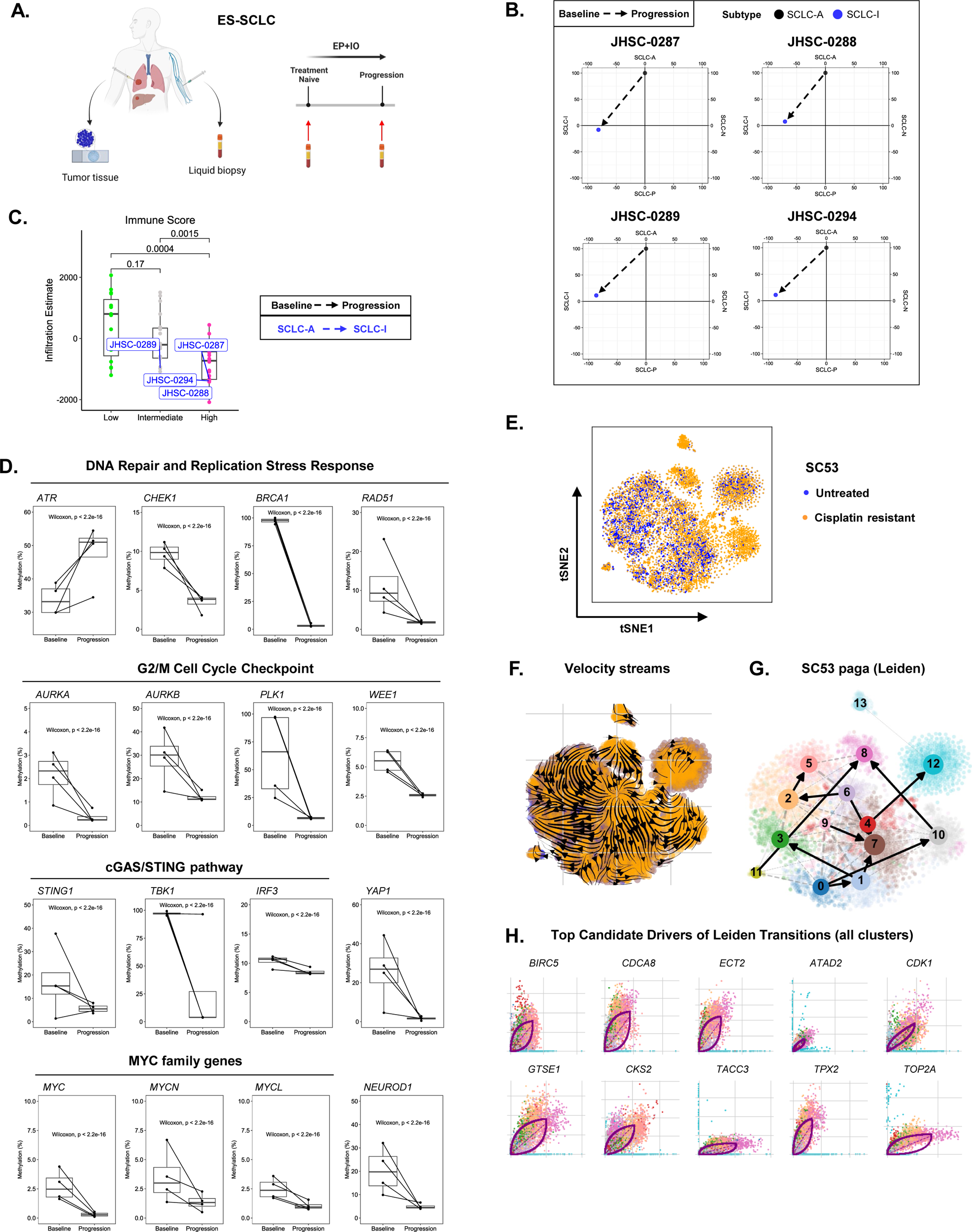
DDR status is linked to subtype switching and unique routes of disease progression following frontline chemoimmunotherapy. **A.** ES-SCLC patient matched tissue and longitudinal liquid biopsy collection schema. **B.** Subtype switching plots following frontline chemoimmunotherapy. **C.** Immune cell infiltration levels in diagnostic tissue samples from subtype switching patients shown in (**B**). **D.** Promoter methylation changes in subtype switching patients from baseline to progression on frontline chemoimmunotherapy. **E.** SC53 untreated and cisplatin relapsed scRNAseq tSNE. **F.** SC53 untreated and cisplatin relapsed RNA velocity streams. **G.** SC53 untreated and cisplatin relapsed Leiden clusters. **H.** Top candidate drivers of cell transitions between Leiden clusters.

To better understand potential mechanisms driving these plastic tumors, we analyzed ctDNA methylation data from matched treatment naïve and progression liquid biopsies. This analysis demonstrated that subtype switching tumors exhibit bifurcated phenotypes enforcing both 1) DNA repair and cell cycle arrest and 2) cGAS-STING, YAP1 related inflammatory programs (**Figure 6D**). Specifically, we observed substantial decreases in promoter methylation of genes mediating DNA repair and replication stress responses, translesion synthesis polymerases, G2/M cell cycle arrest, and cGAS-STING signaling (**Figure 6D, Supplementary Figure 14**). We also found decreased methylation of *YAP1*, reminiscent of previous studies demonstrating a *MYC-NEUROD1-YAP1* axis driving plasticity in SCLC (**Figure 6D**) [45]. Interestingly, we did not find decreased methylation of immune inhibitory checkpoints that could also promote tumor cell survival despite acquiring ‘inflamed’ features (**Supplementary Figure 14**). These data are consistent with chemotherapy induced DNA damage driving both G2/M cell cycle arrest and a cGAS-STING mediated inflammatory signaling response in SCLC tumor cells.

To validate this model, we analyzed scRNAseq data from DDR Intermediate PDX model SC53 at both cisplatin sensitive and resistant timepoints (**Figure 6E**). As in human tumors, cells with “inflamed” features emerge following platinum resistance [5]. Using RNA velocity analyses, we found that genes regulating G2/M cell cycle arrest, replication stress, and DNA repair are top drivers of cell plasticity in this model (**Figure 6F-H, Supplementary Figures 15-16, Supplementary Table 2**). Additionally, we identified genes upregulated by cGAS/STING signaling as additional drivers of plasticity (**Supplementary Table 2**) [46]. While multiple cell states exist in both untreated and cisplatin resistant tumors, random walk simulations demonstrated an overwhelming convergence on Leiden cluster 8, which is characterized by genes regulating G2/M cell cycle arrest and DNA repair—phenocopying our longitudinal liquid biopsy results (**Supplementary Figure 14, Supplementary Figure 16-17**) [36]. Together, these data link baseline DDR status to subtype switching and unique routes of disease progression following frontline therapy.

## Discussion

Small cell lung cancer is a highly aggressive malignancy characterized by recalcitrance to therapy and rapid progression to heterogeneous phenotypes following treatment. In this study, we demonstrate that SCLC tumors cluster into three DDR phenotypes with unique molecular features not captured by subtype assignments alone. Our data show that initial DDR status is linked to chemotherapy response, tumor immune evasion, and evolution to plastic phenotypes following frontline therapy.

Using multi-omic profiling of human SCLC tumor samples, *in vitro* and *in vivo* model systems, we demonstrate that hallmarks of increasing DDR status include increased expression of DNA repair and cell cycle checkpoint effectors, elevated levels of replication stress, and heightened G2/M cell cycle arrest. These data are consistent with a model wherein DDR status identifies differing levels of intrinsic DNA damage and replication stress in treatment naïve SCLC tumors. Despite these striking differences across DDR clusters, we did not find an association with SLFN11 expression, which has been previously reported to positively correlate with DDR gene expression. In fact, our results demonstrate that SLFN11 expression is bimodal within DDR clusters, with each cluster containing both SLFN11 High and Low tumors. These data highlights that our DDR clusters identify SCLCs with similar global DNA repair phenotypes not currently captured by SLFN11 expression alone. Clinically, SLFN11 IHC is being used to try and stratify patients more likely to benefit from DNA damaging targeted therapies [47]. Moving forward, it will be important to consider SLFN11 expression in the context of global DDR status, as it stands to reason that combined assessment of bypass pathways (i.e. translesion synthesis) can further enhance the predictive value of SLFN11 expression.

Beyond DNA damage phenotypes, our results show that increased DDR status is linked with decreased “inflamed” features, both within and across SCLC subtypes. Specifically, we find SCLCs with increased DDR exhibit multiple alterations promoting immune evasion including MHC Class I silencing, decreased pro-inflammatory cytokine expression, and cGAS/STING dysregulation. These alterations are coincident with immune cold microenvironments, as evidenced by decreased immune cell infiltration signatures observed in two independent patient cohorts. Importantly, we demonstrate that these results are independent of the inflamed subtype, whereas stratifying only SCLC-A tumors by their DDR status identifies a spectrum of immune cell infiltration signatures and inflamed biomarker prevalence. Additionally, we establish that these observations are independent of tumor purity by recapitulating our results in pure tumor cell populations both *in vitro* and *in vivo*. It should be noted that DDR specific immune evasive phenotypes are tightly correlated with LSD1 protein expression. Our findings recapitulate reports from several groups linking elevated LSD1 expression with MHC Class I silencing in SCLC [6, 42, 43]. Additionally, our data contextualizes and further extends these findings by linking upregulated LSD1 with elevated replication stress and increased DDR signature expression. Collectively, these data support an intriguing model where tumors with increased levels of replication stress are dependent on LSD1-mediated MHC Class I silencing and immune evasion for tumor development and maintenance. Based on these findings, we hypothesize that tumors with elevated DDR may exhibit increased responses to combined LSD1 inhibitor and immunotherapy combinations currently being evaluated in SCLC (NCT05191797). Moving forward, it will be crucial to test whether targeting other immune checkpoints—such as CTLA4, CD276, or LAG3—can better control disease progression in seemingly more immune evasive DDR Intermediate and High tumors.

To our knowledge, our study represents one of the largest analyses of transcriptional markers and frontline chemotherapy outcomes in ES-SCLC. Our work suggests that treatment naïve DDR status identifies SCLC tumors with differing depth and duration of response to frontline chemotherapy. These results provide insight into the well-recognized paradox of SCLC’s initial exquisite chemotherapy sensitivity and propensity for rapid chemoresistance [1]. Together, our results support a model where high levels of replication stress and cell-intrinsic DNA damage drive compensatory upregulation of the DDR machinery in treatment naïve tumors, priming SCLCs for resistance to DNA damaging therapies and potentially leading to poor patient outcomes. Our finding that DDR High and Intermediate patients have numerically shortened overall survival following frontline chemotherapy, compared to DDR Low, could be further explained by the fact that many additional lines of therapy in SCLC are also DNA-damage based. Thus, as these tumors quickly develop resistance to frontline DNA-damaging therapies, they are further primed for cross-resistance to additional lines of therapy with similar mechanisms of action, a phenomena recently confirmed in SCLC [48]. Interestingly, multi-omic profiling of preclinical SCLC models demonstrated that tumors with increased DDR signatures exhibited elevated expression of the epigenetic regulator SMYD3. Recently, SMYD3 has been shown to drive resistance to alkylating chemotherapeutic agents and that pharmacologic inhibition of SMYD3 reverses resistance in SCLC models [49]. Moving forward, it will be intriguing to test if targeting SMYD3 can increase efficacy of second line lurbinectedin in a DDR specific fashion.

Lastly, our data highlight that treatment naïve DDR status has implications for SCLC evolution. Using longitudinal liquid biopsies, we find that tumors with elevated DDR become highly plastic and can acquire non-neuroendocrine features following frontline therapy. Paradoxically, these plastic tumors shift subtypes towards a more “inflamed” state but do not derive durable benefit from frontline chemoimmunotherapy targeting the PD1 axis. Additionally, these plastic tumors have some of the most immune cell poor tumor microenvironments at baseline. Mechanistically, we find these tumors upregulate bifurcated phenotypes regulating both DNA repair and inflammatory programs. These data are reminiscent of findings from Lissa et al where SCLCs with hybrid neuroendocrine and non-neuroendocrine features respond poorly to therapy and rapidly progress [12]. Interestingly, in their report the authors find that these hybrid tumors are resistant to targeted ATR inhibition. In our study, we find that plastic tumors with similar features converge on a G2/M arrest and increasingly methylate *ATR*, providing a potential explanation for the observed lack of response to ATR inhibition. Our results instead argue for targeting these hybrid tumors with agents that inhibit G2/M cell cycle effectors, such as PLK1, AURK, and WEE1 [50]. Furthermore, we find that these plastic tumors upregulate the *MYC*/*NEUROD1*/*YAP1* axis previously shown to drive plasticity in SCLC [45]. Our data extend previous reports by linking *YAP1* upregulation with coincident *cGAS-STING* and inflammatory signaling activity following therapy, providing a potential mechanism for *YAP1* upregulation. We also observed an upregulation of several TLS polymerases at progression in these plastic tumors. Recent work has demonstrated that upregulation of the TLS machinery underpins resistance to a host of DNA damaging agents in SCLC [48]. Our work provides further rationale for exploring targeted TLS inhibitor combinations in relapsed tumors as a mechanism for chemosensitization.

Our study has several limitations. First, our study is retrospective in nature. Second, analysis of IMPower133 patient outcomes following frontline chemotherapy were underpowered for biomarker subgroup analyses. Third, we lacked matched genomic profiling data from our human patient cohorts and thus could not determine the underlying genetic drivers of distinct DDR phenotypes. Lastly, our analysis of DDR specific subtype switching is constrained by a small sample size. Future studies with larger cohorts will be needed to confirm our findings.

In conclusion, we establish that SCLC clusters into three biologically distinct, clinically relevant DDR clusters. Our work demonstrates that initial DDR status plays a key role in shaping SCLC phenotypes, chemotherapy response, and tumor evolution following frontline therapy. Future work targeting DDR cluster specific phenotypes will be instrumental for ultimately improving SCLC patient outcomes.

## Supporting information

Morris SCLC DDR Supplementary Tables

## Acknowledgements

This work was supported by: NIH/NCI CCSG P30-CA016672 (MD Anderson Bioinformatics Shared Resource, and the MD Anderson Institutional Tissue bank (ITB)); The University of Texas-Southwestern and MD Anderson Cancer Center Lung SPORE P50-CA070907 (JW, JVH, CMG, LAB); NIH/NCI R01-CA207295 (LAB); NIH/NCI U01-CA213273 (JVH, LAB); NIH/NCI R50-CA243698 (CAS); NIH/NCI U01-CA256780 (JVH, LAB); NIH/NCI U24-CA213274 (LAB); The Department of Defense LC210510 (LAB); the LUNGevity Foundation 2020-02 (CMG); CPRIT RP210159 (CMG); Andrew Sabin Family Foundation (CMG); NETRF (CMG); NIH/NCI T32-CA009666 (KC); Rexanna’s Foundation for Fighting Lung Cancer (JVH, LAB, CMG); Jeffrey Lee Cousins Endowed Fellowship in Lung Cancer Research (BBM); and generous philanthropic contributions to The University of Texas MD Anderson Lung Cancer Moon Shot Program (JVH, CMG, LAB). We would also like to thank A.R.K., L.W.Y., M.J.A., R.B.N., K.E.N., J.O., J.K.R., B.&B.N, C.K., P.C.B., S.S., and W.A.B. for their philanthropic support of this project. BBM is a TRIUMPH Fellow in the CPRIT Research Training Program (RP210028). This research includes work performed in the Single Cell Genomics Core Facility, which is supported in part by CPRIT Single Core grant RP180684. The authors would like to thank the patients that participated in this study, as well as their families.

## Supplementary Figure Legends

**Supplementary Figure 1:**
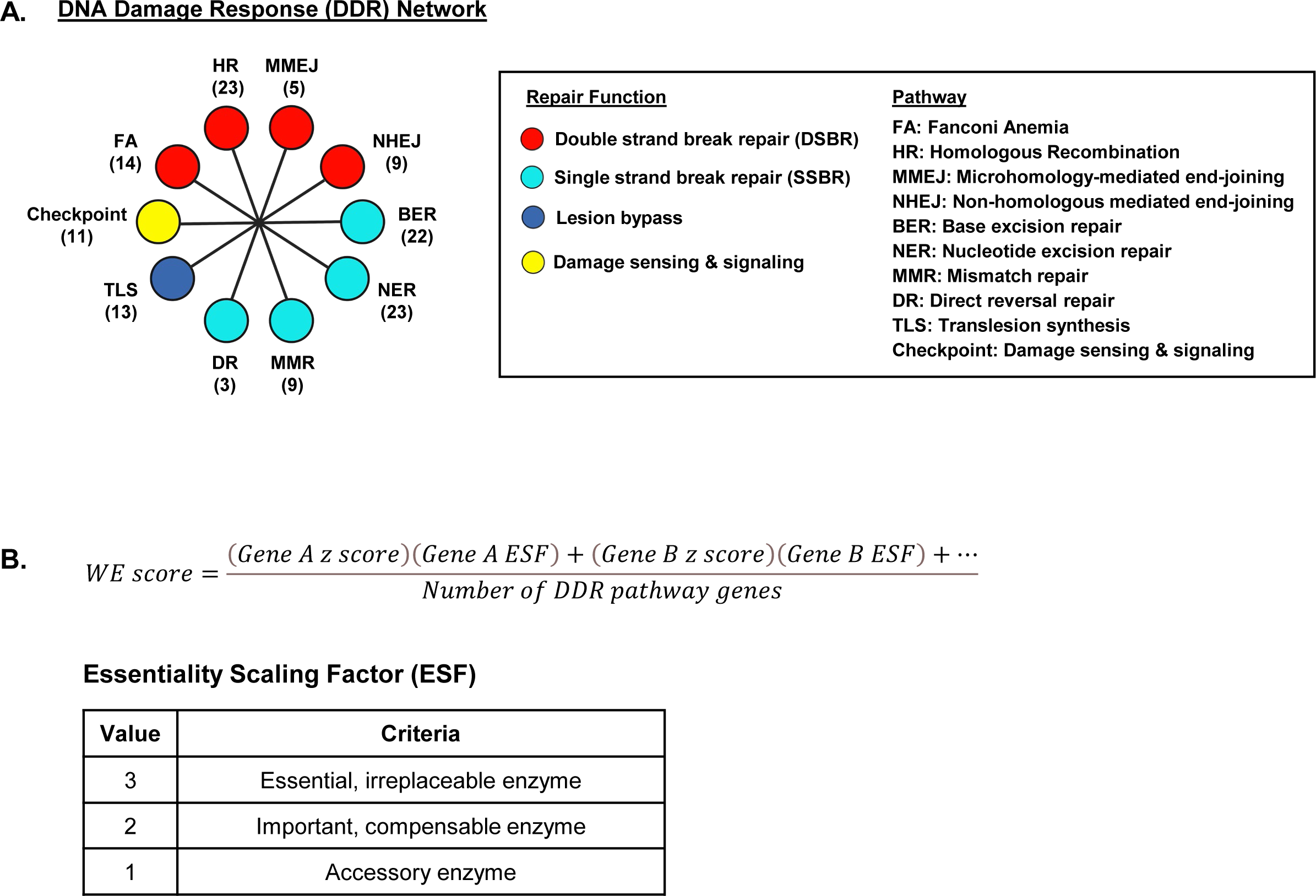
DDR subtyping analysis overview. **A.** DDR network overview. HR: Homologous recombination. MMEJ: microhomology-mediated end-joining. NHEJ: Non-homologous end-joining. BER: Base excision repair. NER: Nucleotide excision repair. MMR: Mismatch repair. DR: Direct reversal repair. TLS: Translesion synthesis. Checkpoint: Damage sensing and signaling. FA: Fanconi Anemia. Numbers in parentheses indicate number of pathway genes analyzed by our method. **B.** WE score formula and Essentiality Scaling Factor (ESF) criteria.

**Supplementary Figure 2:**
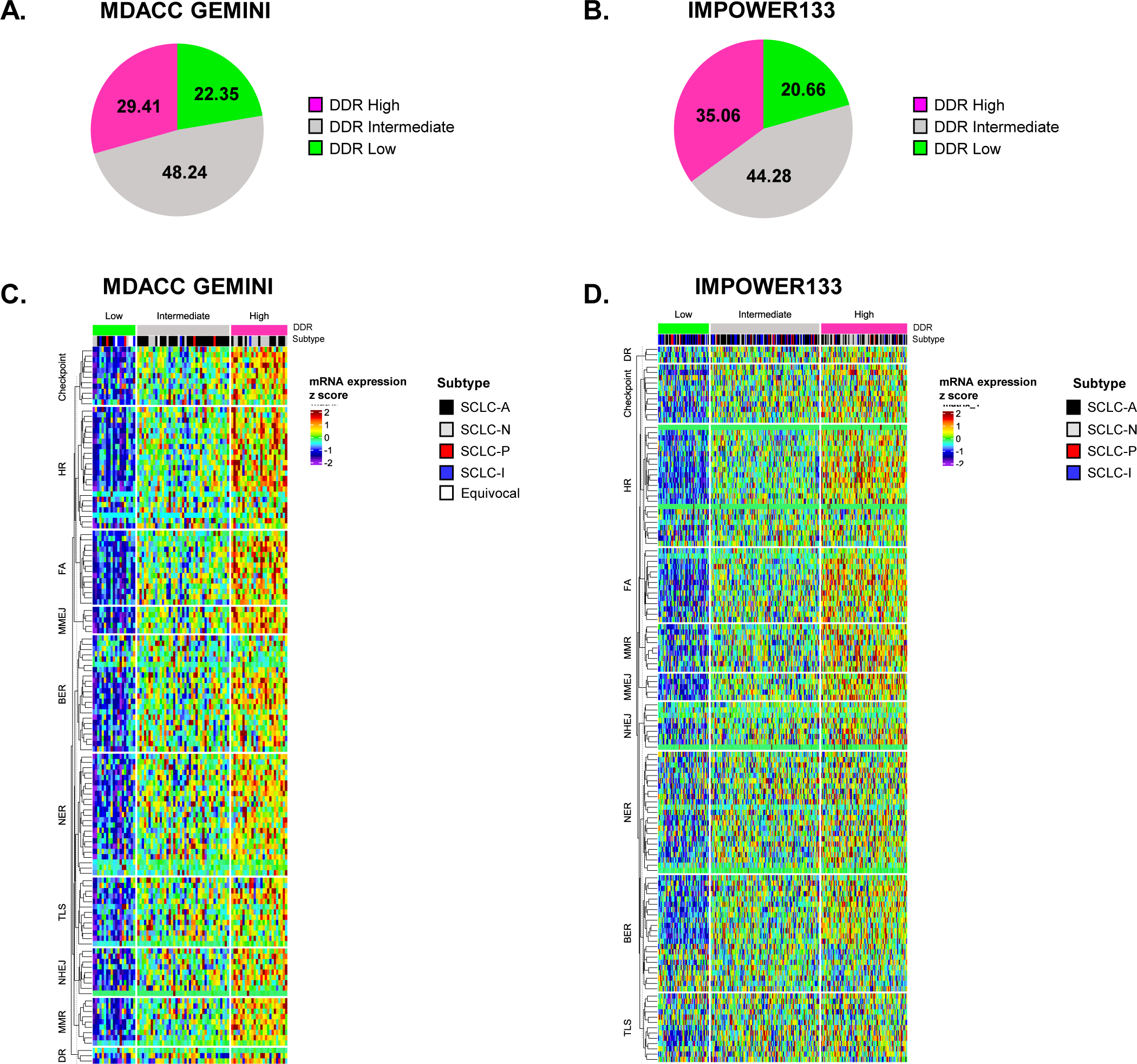
DDR cluster prevalence and DDR pathway single gene heatmaps. **A.** DDR cluster prevalence in GEMINI cohort. **B.** DDR cluster prevalence in IMPOWER133 cohort. **C.** GEMINI DDR pathway single gene expression heatmap. **D.** IMPOWER133 DDR pathway single gene expression heatmap.

**Supplementary Figure 3:**
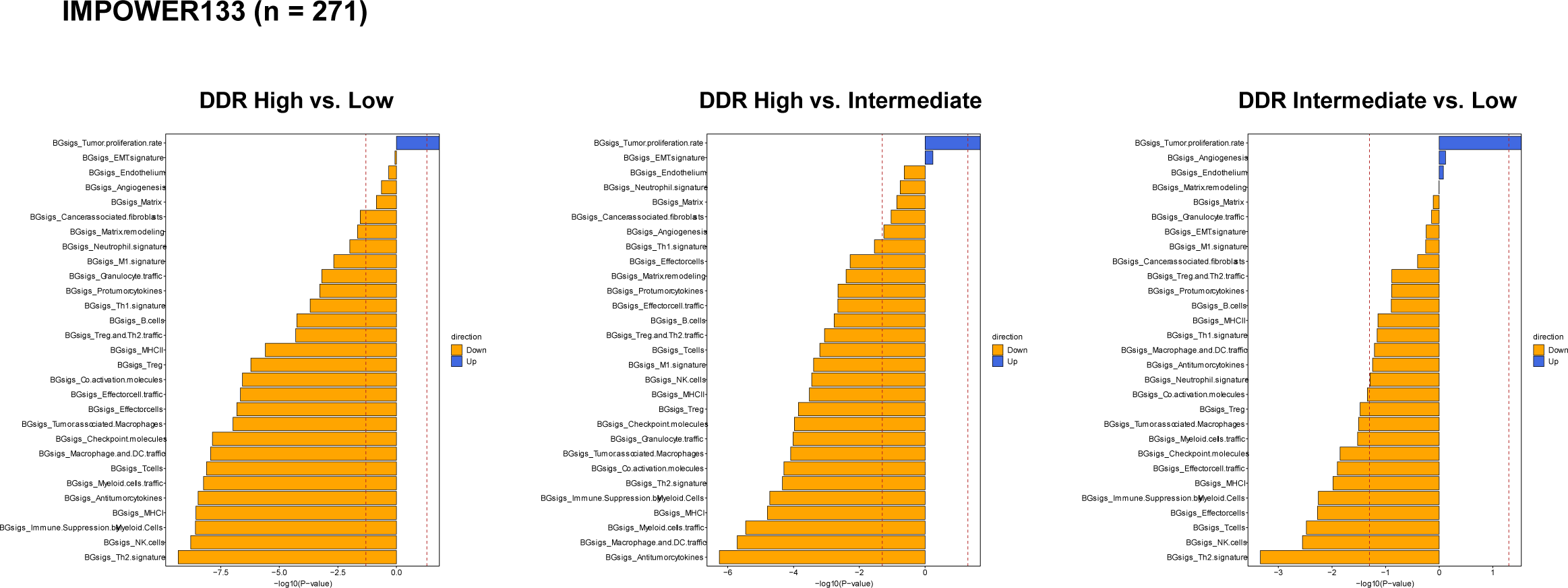
IMpower133 DDR cluster QuSAGE BG signature results.

**Supplementary Figure 4:**
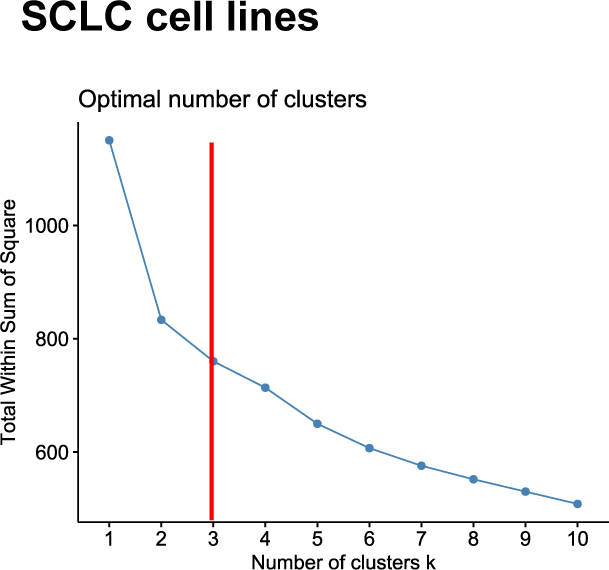
SCLC CellMiner cell line optimal number of k clusters elbow plot.

**Supplementary Figure 5:**
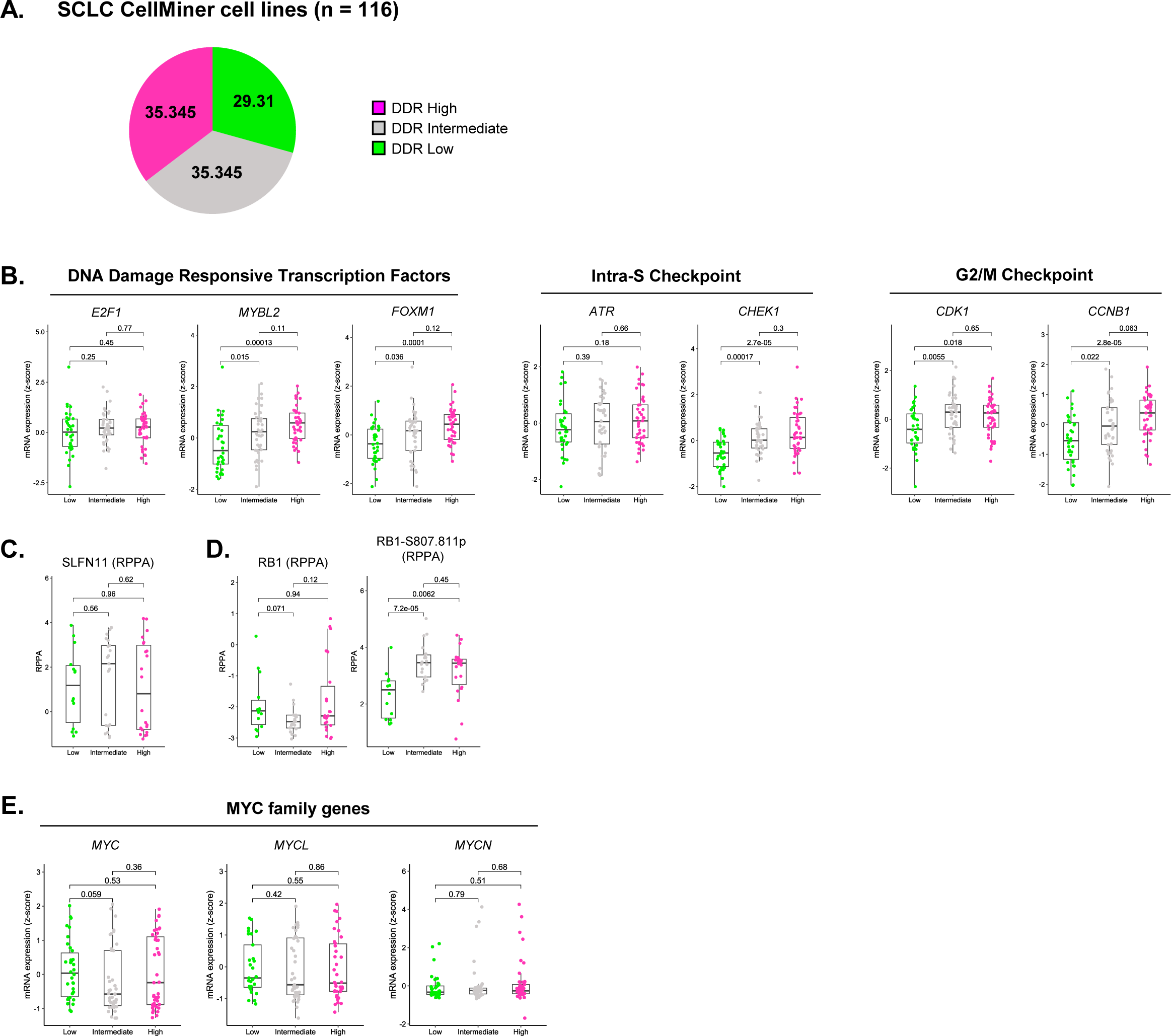
SCLC CellMiner cell line DDR cluster prevalence and marker expression. **A.** DDR cluster prevalence in SCLC cell line models. **B.** DNA damage responsive transcription factors, intra-S cell cycle checkpoint, and G2/M cell cycle checkpoint machinery expression across SCLC cell line DDR clusters**. C.** SLFN11 protein expression in SCLC cell line DDR clusters. **D.** Total RB1 and RB1-S807.S811 phospho protein expression across SCLC cell line DDR clusters. **E.** Cell line DDR cluster MYC family gene expression.

**Supplementary Figure 6:**
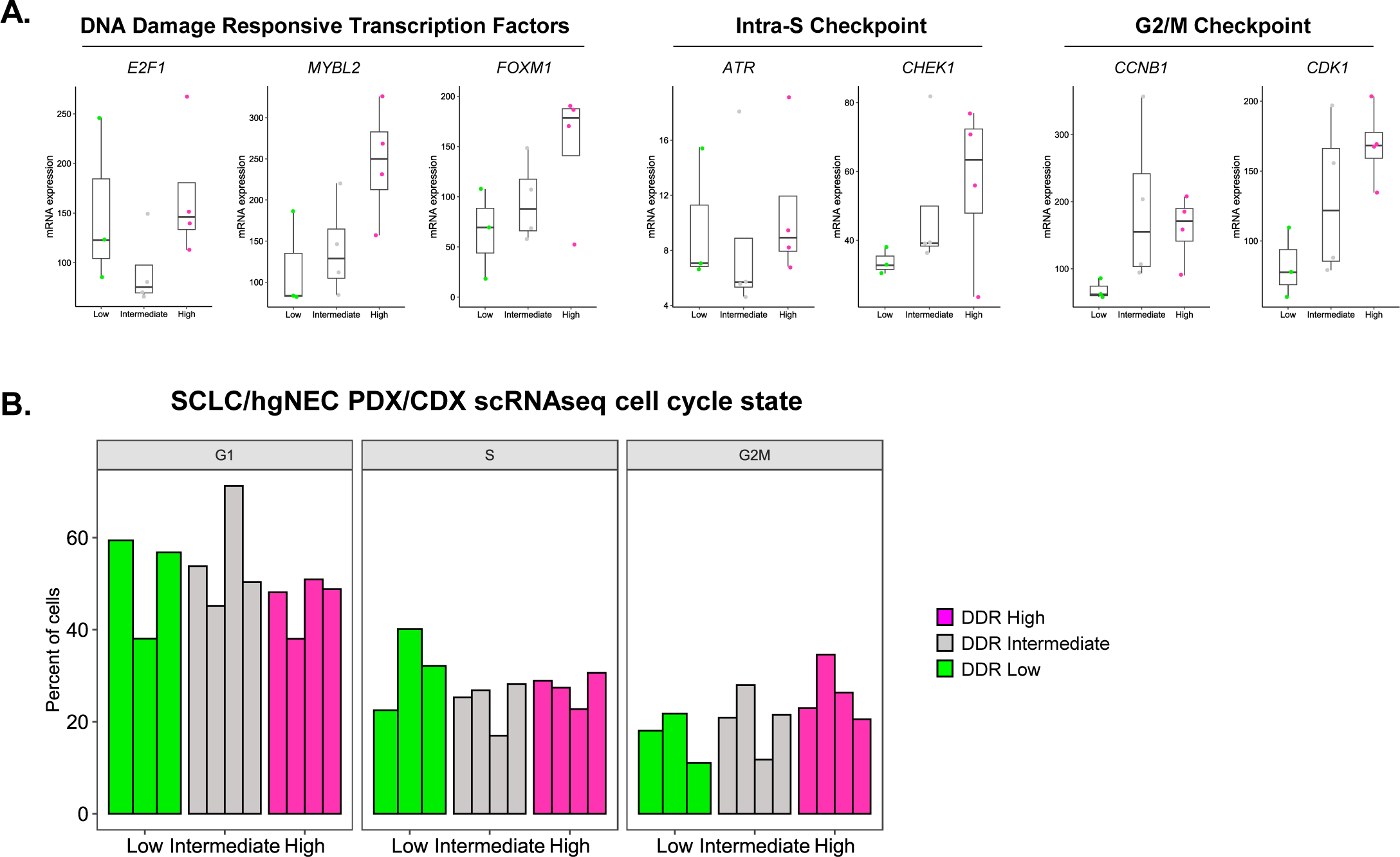
SCLC/hgNEC PDX/CDX DDR gene expression and cell cycle state distribution. **A.** Expression of DNA damage responsive transcription factors, intra-S, and G2/M cell cycle checkpoint effectors in PDX/CDX DDR clusters. **B.** PDX/CDX DDR cluster scRNAseq cell cycle state distributions.

**Supplementary Figure 7:**
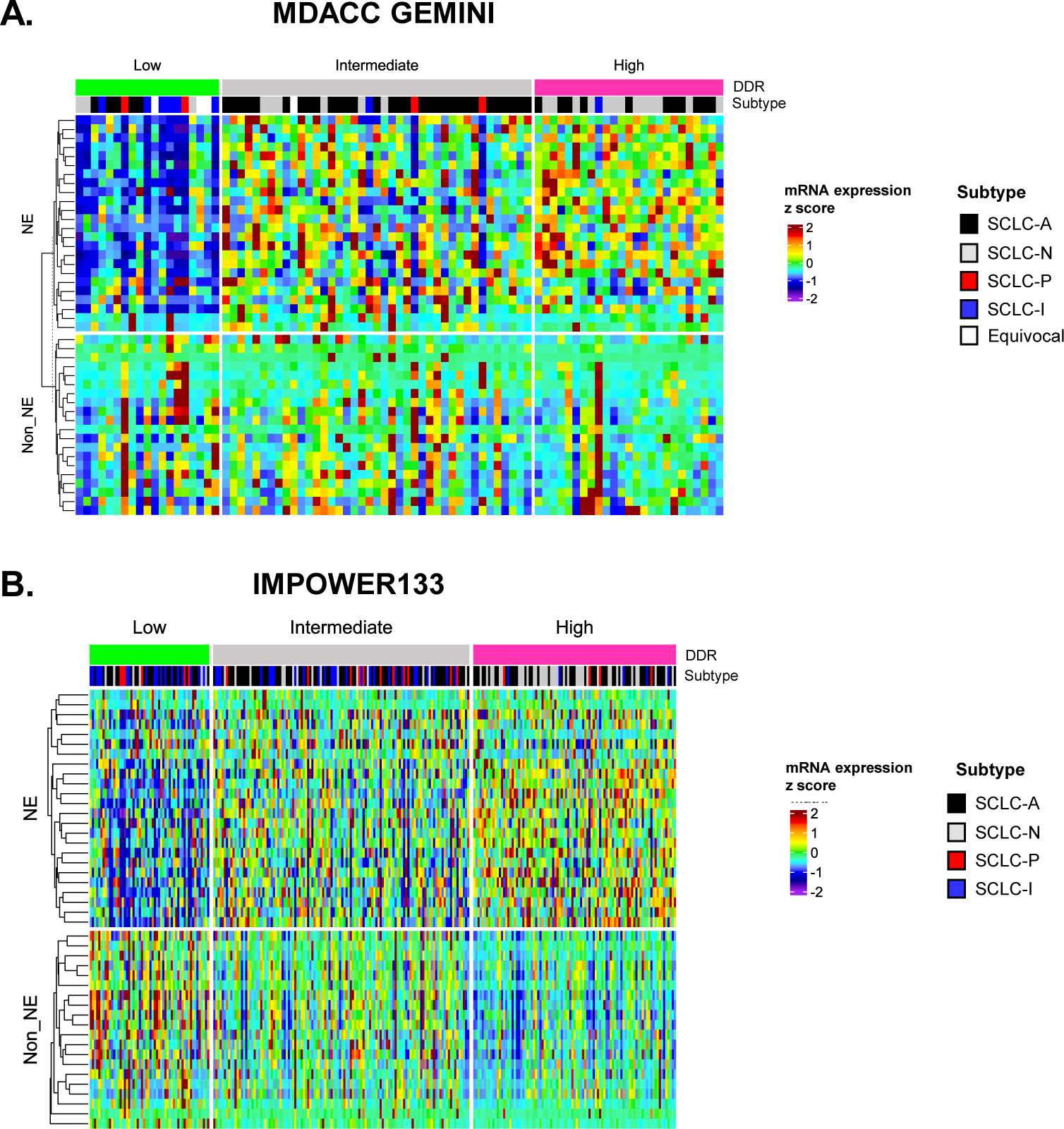
GEMINI and IMPOWER133 DDR cluster neuroendocrine score single gene heatmaps. **A.** GEMINI DDR cluster neuroendocrine score single gene expression heatmap. **B.** IMPOWER133 DDR cluster neuroendocrine score single gene expression heatmap.

**Supplementary Figure 8:**
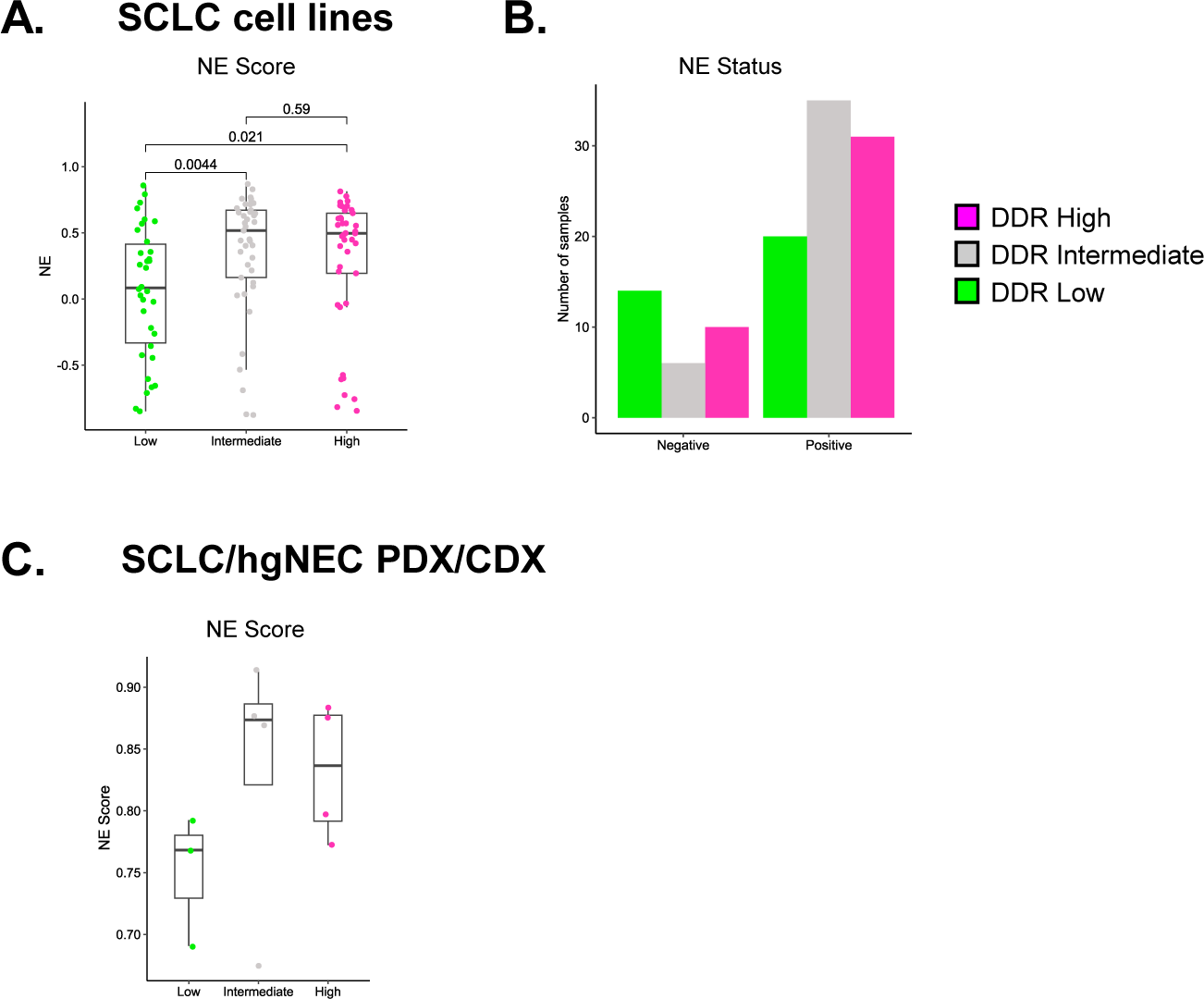
SCLC CellMiner DDR cluster neuroendocrine features. **A.** SCLC cell line DDR cluster NE scores. **B.** SCLC cell line DDR cluster neuroendocrine status as reported by Tlemsani et al. **C.** SCLC PDX/CDX DDR cluster NE scores.

**Supplementary Figure 9:**
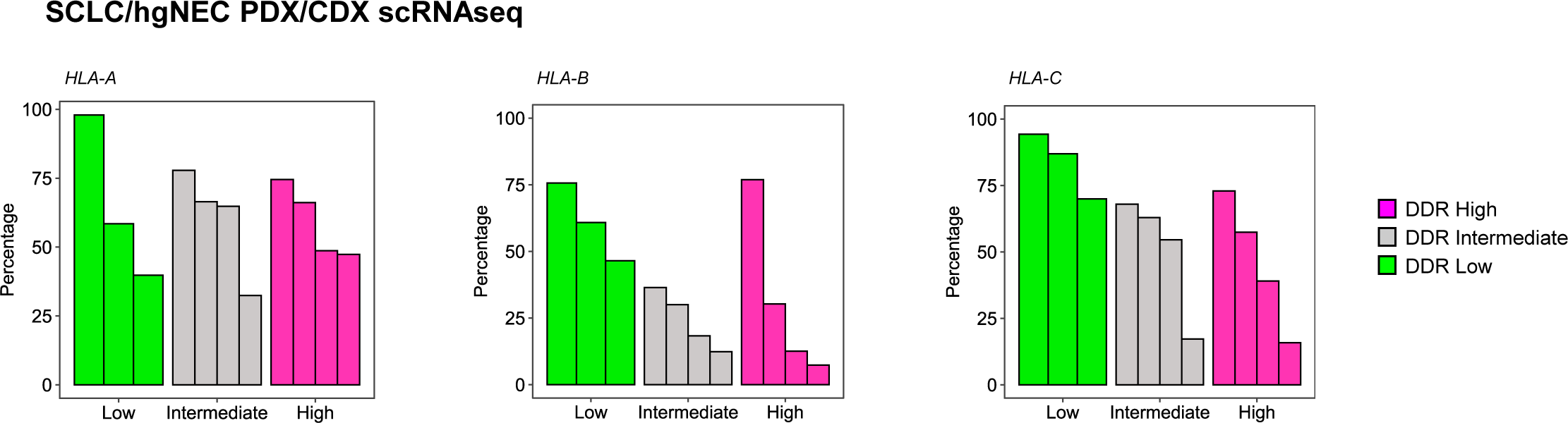
SCLC/hgNEC PDX/CDX DDR cluster MHC Class I scRNAseq expression.

**Supplementary Figure 10:**
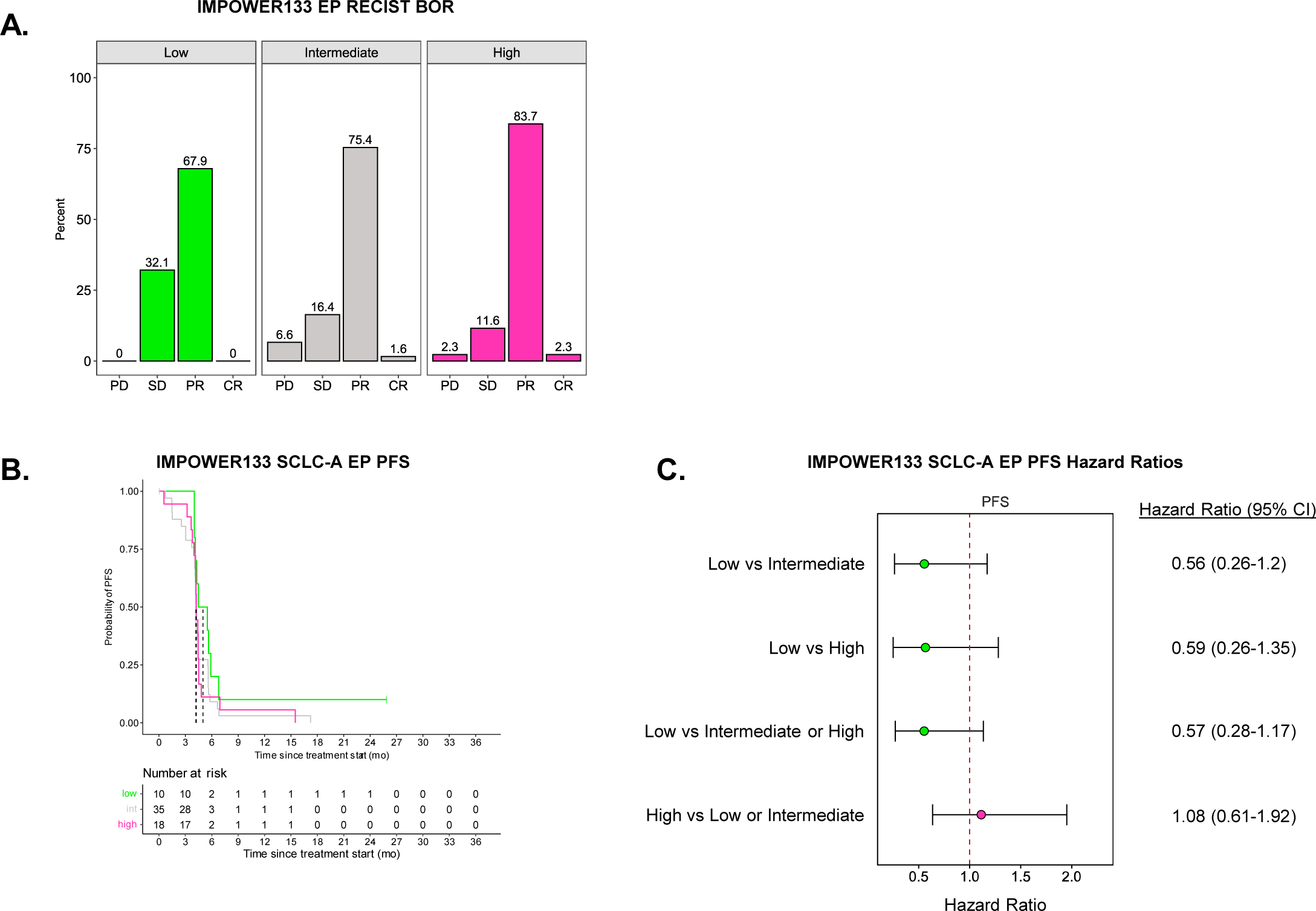
IMPOWER133 DDR cluster chemotherapy response analysis. **A.** RECIST Best Overall Response (BOR) for IMPOWER133 DDR clusters following frontline EP chemotherapy. PD: Progressive disease. SD: Stable disease. PR: Partial response. CR: Complete response. **B.** Progression free survival Kaplan Meier plot for SCLC-A DDR clusters following frontline EP chemotherapy. **C.** Forest plot for progression free survival for SCLC-A DDR clusters following frontline EP chemotherapy.

**Supplementary Figure 11:**
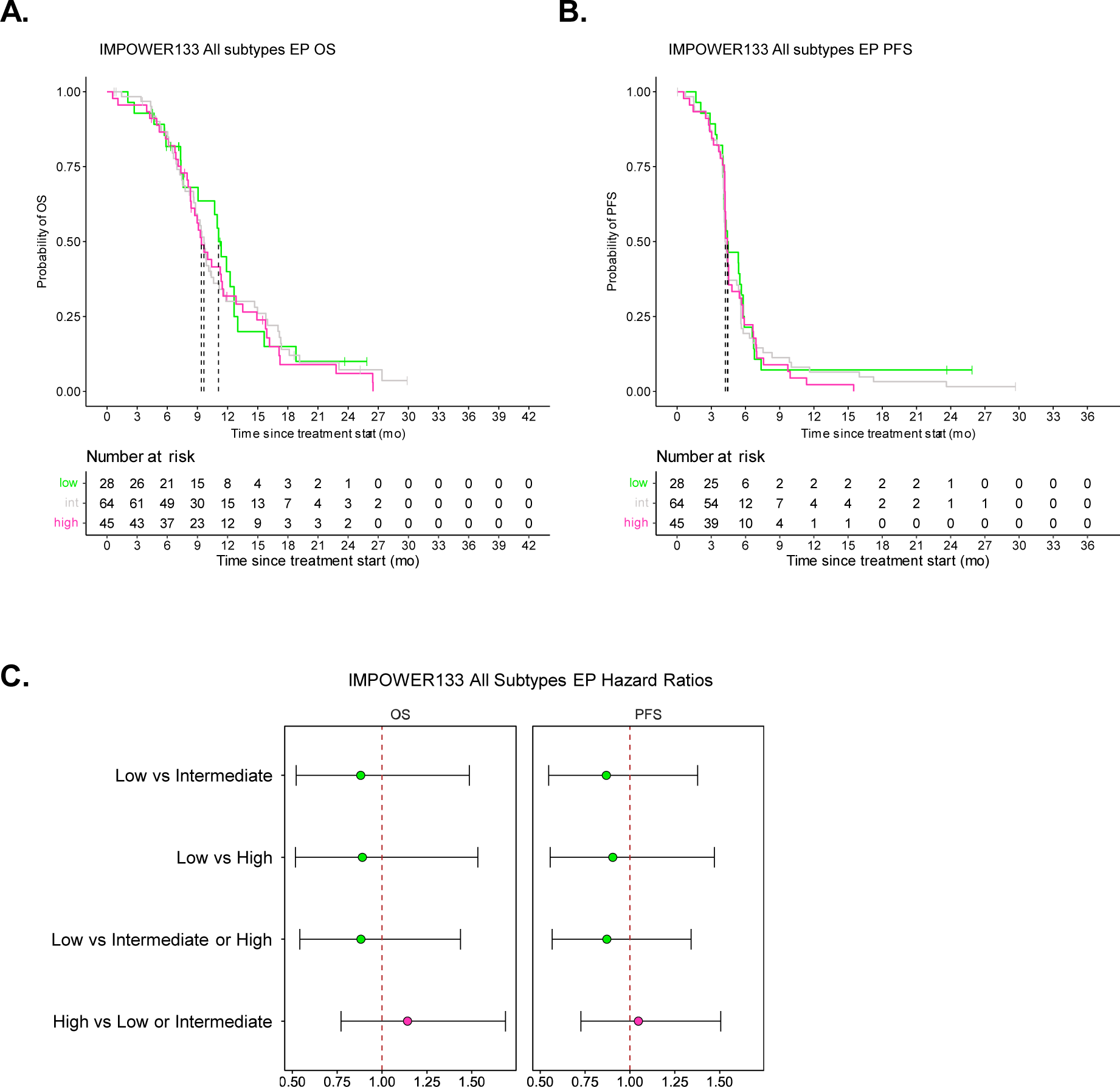
IMPOWER133 DDR cluster all subtypes chemotherapy response analysis. **A.** Overall survival Kaplan Meier plot for all subtypes DDR clusters following frontline EP chemotherapy. **B.** Progression free survival Kaplan Meier plot for all subtypes DDR clusters following frontline EP chemotherapy. **C.** Forrest plots for all subtype DDR cluster OS and PFS outcomes following frontline EP chemotherapy.

**Supplementary Figure 12:**
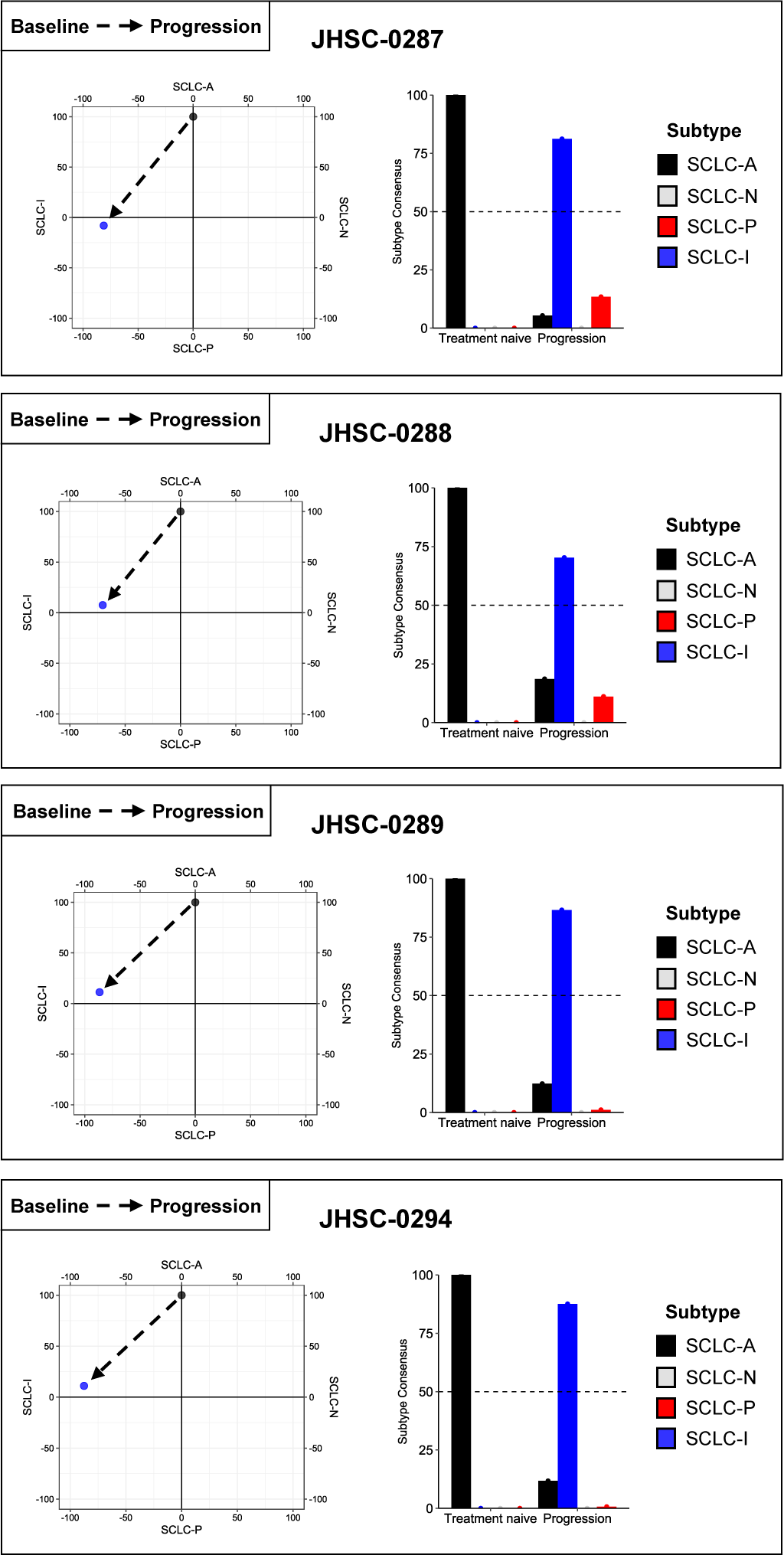
Subtype switching plots. Subtype space and SCLC-DMC subtype calls for subtype switching patients.

**Supplementary Figure 13:**
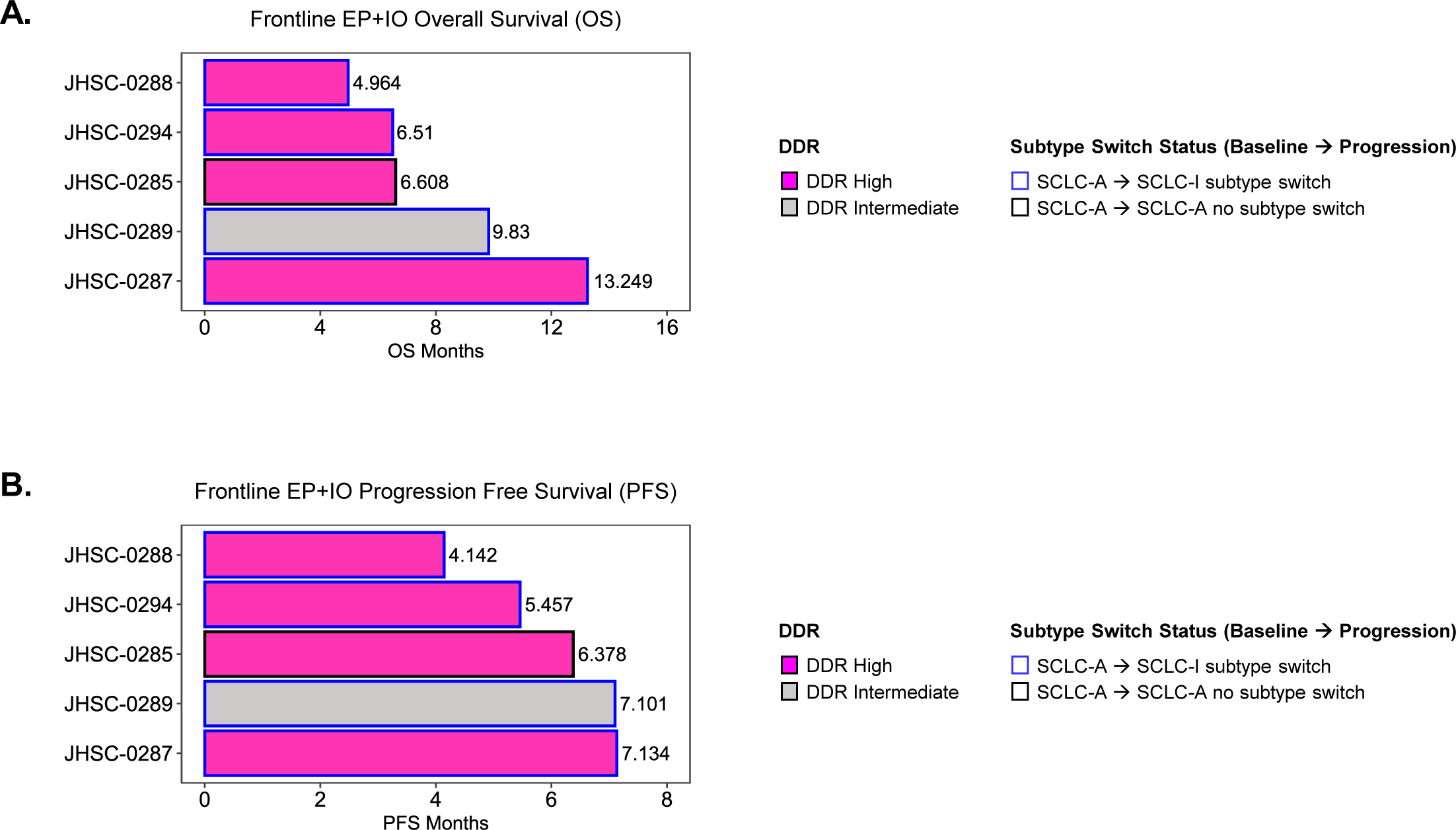
Subtype switching patient outcomes following frontline chemoimmunotherapy. **A.** Overall survival following frontline chemoimmunotherapy. **B.** Progression free survival following frontline chemoimmunotherapy.

**Supplementary Figure 14:**
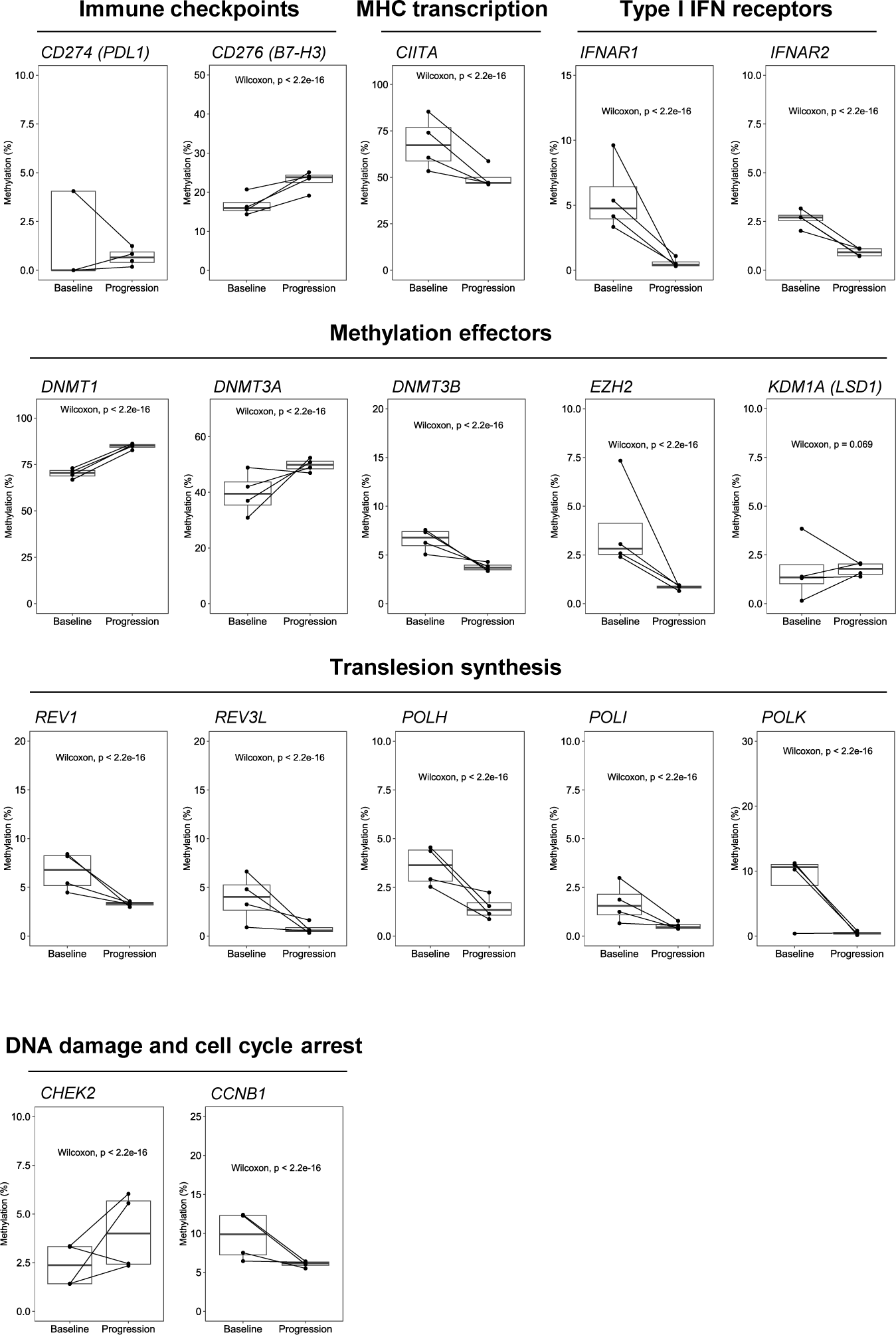
Promoter methylation changes for subtype switching patients from baseline to progression following frontline chemoimmunotherapy.

**Supplementary Figure 15:**
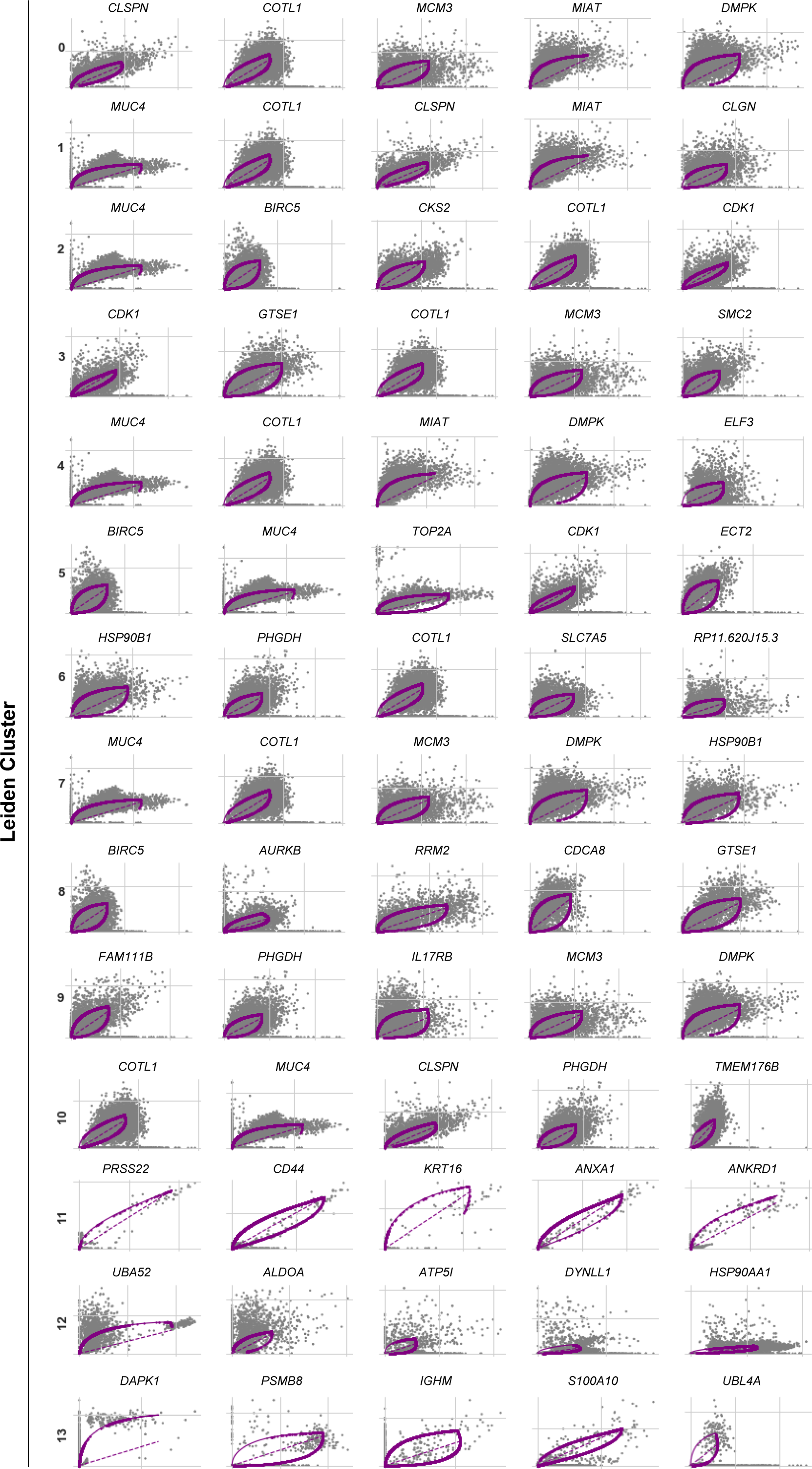
SC53 Leiden cluster specific top drivers of plasticity.

**Supplementary Figure 16:**
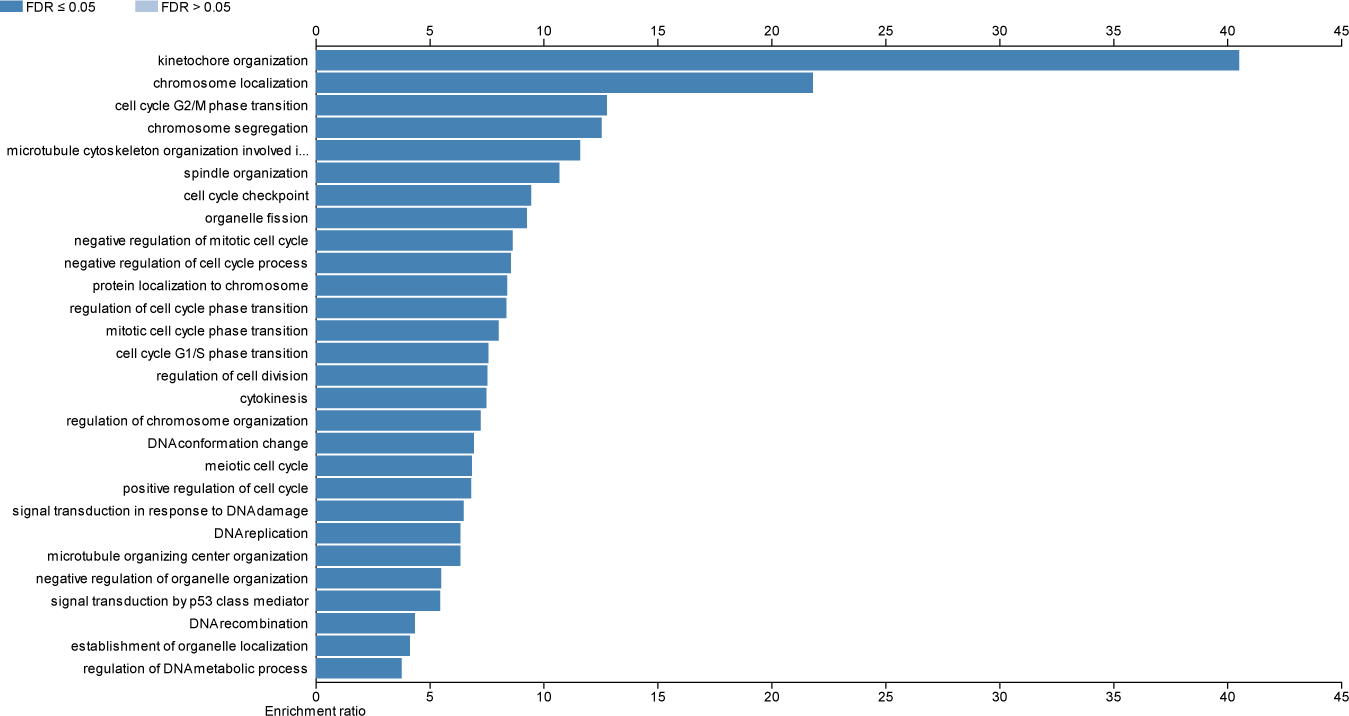
WebGestalt over-representation analysis of Leiden Cluster 8 genes.

**Supplementary Figure 17:**
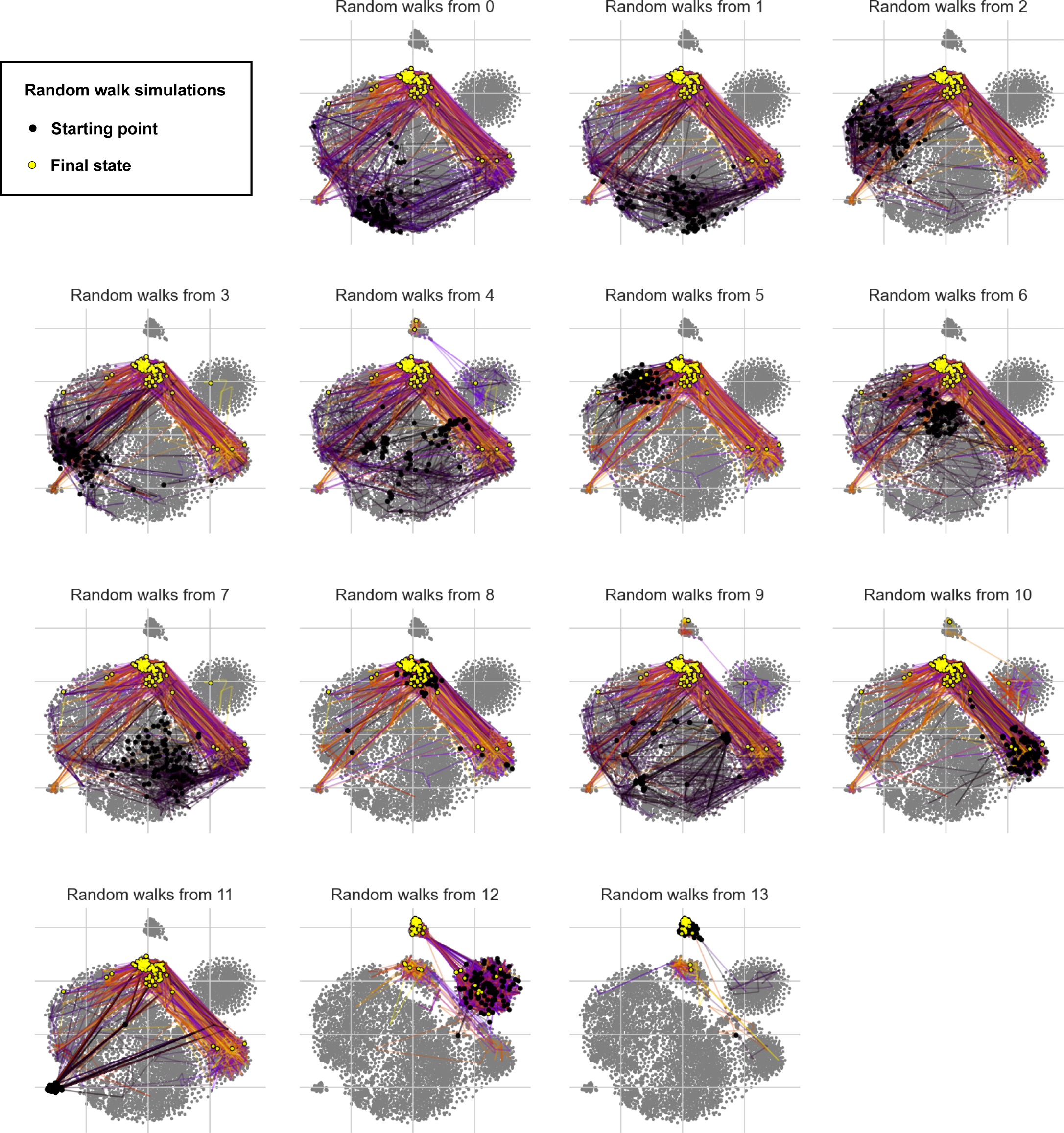
SC53 Leiden cluster specific random walk simulations.

**Supplementary Table 1: WE score individual DDR gene Essentiality Scaling Factor (ESF) assignments and example raw WE score generation.**

**Supplementary Table 2: SC53 dynamic genes by Leiden cluster.**

## References

1. Rudin CM, Brambilla E, Faivre-Finn C, Sage J: Small-cell lung cancer. Nat Rev Dis Primers 2021, 7:3.

2. George J, Lim JS, Jang SJ, Cun Y, Ozretic L, Kong G, Leenders F, Lu X, Fernandez-Cuesta L, Bosco G, et al: Comprehensive genomic profiles of small cell lung cancer. Nature 2015, 524:47–53.

3. Tan AC, Tan DSW: Targeted Therapies for Lung Cancer Patients With Oncogenic Driver Molecular Alterations. J Clin Oncol 2022, 40:611–625.

4. Horn L, Mansfield AS, Szczesna A, Havel L, Krzakowski M, Hochmair MJ, Huemer F, Losonczy G, Johnson ML, Nishio M, et al: First-Line Atezolizumab plus Chemotherapy in Extensive-Stage Small-Cell Lung Cancer. N Engl J Med 2018, 379:2220–2229.

5. Gay CM, Stewart CA, Park EM, Diao L, Groves SM, Heeke S, Nabet BY, Fujimoto J, Solis LM, Lu W, et al: Patterns of transcription factor programs and immune pathway activation define four major subtypes of SCLC with distinct therapeutic vulnerabilities. Cancer Cell 2021, 39:346–360 e347.

6. Rudin CM, Balli D, Lai WV, Richards AL, Nguyen E, Egger JV, Choudhury NJ, Sen T, Chow A, Poirier JT, et al: Clinical Benefit From Immunotherapy in Patients With SCLC Is Associated With Tumor Capacity for Antigen Presentation. J Thorac Oncol 2023.

7. Nabet BY, Hamidi H, Lee MC, Banchereau R, Morris S, Adler L, Gayevskiy V, Elhossiny AM, Srivastava MK, Patil NS, et al: Immune heterogeneity in small-cell lung cancer and vulnerability to immune checkpoint blockade. Cancer Cell 2024, 42:429–443 e424.

8. Morris BB, Smith JP, Zhang Q, Jiang Z, Hampton OA, Churchman ML, Arnold SM, Owen DH, Gray JE, Dillon PM, et al: Replicative Instability Drives Cancer Progression. Biomolecules 2022, 12.

9. Huang D, Savage SR, Calinawan AP, Lin C, Zhang B, Wang P, Starr TK, Birrer MJ, Paulovich AG: A highly annotated database of genes associated with platinum resistance in cancer. Oncogene 2021, 40:6395–6405.

10. Hoppe MM, Jaynes P, Wardyn JD, Upadhyayula SS, Tan TZ, Lie S, Lim DGZ, Pang BNK, Lim S, J PSY, et al: Quantitative imaging of RAD51 expression as a marker of platinum resistance in ovarian cancer. EMBO Mol Med 2021, 13:e13366.

11. Takahashi N, Kim S, Schultz CW, Rajapakse VN, Zhang Y, Redon CE, Fu H, Pongor L, Kumar S, Pommier Y, et al: Replication stress defines distinct molecular subtypes across cancers. Cancer Res Commun 2022, 2:503–517.

12. Lissa D, Takahashi N, Desai P, Manukyan I, Schultz CW, Rajapakse V, Velez MJ, Mulford D, Roper N, Nichols S, et al: Heterogeneity of neuroendocrine transcriptional states in metastatic small cell lung cancers and patient-derived models. Nat Commun 2022, 13:2023.

13. Heeke S, Gay CM, Estecio MR, Tran H, Morris BB, Zhang B, Tang X, Raso MG, Rocha P, Lai S, et al: Tumor- and circulating-free DNA methylation identifies clinically relevant small cell lung cancer subtypes. Cancer Cell 2024.

14. Tlemsani C, Pongor L, Elloumi F, Girard L, Huffman KE, Roper N, Varma S, Luna A, Rajapakse VN, Sebastian R, et al: SCLC-CellMiner: A Resource for Small Cell Lung Cancer Cell Line Genomics and Pharmacology Based on Genomic Signatures. Cell Rep 2020, 33:108296.

15. Byers LA, Wang J, Nilsson MB, Fujimoto J, Saintigny P, Yordy J, Giri U, Peyton M, Fan YH, Diao L, et al: Proteomic profiling identifies dysregulated pathways in small cell lung cancer and novel therapeutic targets including PARP1. Cancer Discov 2012, 2:798–811.

16. Black SJ, Ozdemir AY, Kashkina E, Kent T, Rusanov T, Ristic D, Shin Y, Suma A, Hoang T, Chandramouly G, et al: Molecular basis of microhomology-mediated end-joining by purified full-length Poltheta. Nat Commun 2019, 10:4423.

17. Ceccaldi R, Sarangi P, D’Andrea AD: The Fanconi anaemia pathway: new players and new functions. Nat Rev Mol Cell Biol 2016, 17:337–349.

18. Jiricny J: The multifaceted mismatch-repair system. Nat Rev Mol Cell Biol 2006, 7:335–346.

19. Krejci L, Altmannova V, Spirek M, Zhao X: Homologous recombination and its regulation. Nucleic Acids Res 2012, 40:5795–5818.

20. Krokan HE, Bjoras M: Base excision repair. Cold Spring Harb Perspect Biol 2013, 5:a012583.

21. Lanz MC, Dibitetto D, Smolka MB: DNA damage kinase signaling: checkpoint and repair at 30 years. EMBO J 2019, 38:e101801.

22. Scharer OD: Nucleotide excision repair in eukaryotes. Cold Spring Harb Perspect Biol 2013, 5:a012609.

23. Stinson BM, Loparo JJ: Repair of DNA Double-Strand Breaks by the Nonhomologous End Joining Pathway. Annu Rev Biochem 2021, 90:137–164.

24. Yang W, Gao Y: Translesion and Repair DNA Polymerases: Diverse Structure and Mechanism. Annu Rev Biochem 2018, 87:239–261.

25. Yi C, He C: DNA repair by reversal of DNA damage. Cold Spring Harb Perspect Biol 2013, 5:a012575.

26. Ritchie ME, Phipson B, Wu D, Hu Y, Law CW, Shi W, Smyth GK: limma powers differential expression analyses for RNA-sequencing and microarray studies. Nucleic Acids Res 2015, 43:e47.

27. Yaari G, Bolen CR, Thakar J, Kleinstein SH: Quantitative set analysis for gene expression: a method to quantify gene set differential expression including gene-gene correlations. Nucleic Acids Res 2013, 41:e170.

28. Yoshihara K, Shahmoradgoli M, Martinez E, Vegesna R, Kim H, Torres-Garcia W, Trevino V, Shen H, Laird PW, Levine DA, et al: Inferring tumour purity and stromal and immune cell admixture from expression data. Nat Commun 2013, 4:2612.

29. Zhang W, Girard L, Zhang YA, Haruki T, Papari-Zareei M, Stastny V, Ghayee HK, Pacak K, Oliver TG, Minna JD, Gazdar AF: Small cell lung cancer tumors and preclinical models display heterogeneity of neuroendocrine phenotypes. Transl Lung Cancer Res 2018, 7:32–49.

30. Stewart CA, Gay CM, Xi Y, Sivajothi S, Sivakamasundari V, Fujimoto J, Bolisetty M, Hartsfield PM, Balasubramaniyan V, Chalishazar MD, et al: Single-cell analyses reveal increased intratumoral heterogeneity after the onset of therapy resistance in small-cell lung cancer. Nat Cancer 2020, 1:423–436.

31. Zheng GX, Terry JM, Belgrader P, Ryvkin P, Bent ZW, Wilson R, Ziraldo SB, Wheeler TD, McDermott GP, Zhu J, et al: Massively parallel digital transcriptional profiling of single cells. Nat Commun 2017, 8:14049.

32. Stuart T, Butler A, Hoffman P, Hafemeister C, Papalexi E, Mauck WM, 3rd, Hao Y, Stoeckius M, Smibert P, Satija R: Comprehensive Integration of Single-Cell Data. Cell 2019, 177:1888–1902 e1821.

33. Bergen V, Lange M, Peidli S, Wolf FA, Theis FJ: Generalizing RNA velocity to transient cell states through dynamical modeling. Nat Biotechnol 2020, 38:1408–1414.

34. Gayoso A, Weiler P, Lotfollahi M, Klein D, Hong J, Streets A, Theis FJ, Yosef N: Deep generative modeling of transcriptional dynamics for RNA velocity analysis in single cells. Nat Methods 2024, 21:50–59.

35. Lange M, Bergen V, Klein M, Setty M, Reuter B, Bakhti M, Lickert H, Ansari M, Schniering J, Schiller HB, et al: CellRank for directed single-cell fate mapping. Nat Methods 2022, 19:159–170.

36. Liao Y, Wang J, Jaehnig EJ, Shi Z, Zhang B: WebGestalt 2019: gene set analysis toolkit with revamped UIs and APIs. Nucleic Acids Res 2019, 47:W199–W205.

37. Carney DN, Gazdar AF, Bepler G, Guccion JG, Marangos PJ, Moody TW, Zweig MH, Minna JD: Establishment and identification of small cell lung cancer cell lines having classic and variant features. Cancer Res 1985, 45:2913–2923.

38. Rudin CM, Poirier JT, Byers LA, Dive C, Dowlati A, George J, Heymach JV, Johnson JE, Lehman JM, MacPherson D, et al: Molecular subtypes of small cell lung cancer: a synthesis of human and mouse model data. Nat Rev Cancer 2019, 19:289–297.

39. Chan JM, Quintanal-Villalonga A, Gao VR, Xie Y, Allaj V, Chaudhary O, Masilionis I, Egger J, Chow A, Walle T, et al: Signatures of plasticity, metastasis, and immunosuppression in an atlas of human small cell lung cancer. Cancer Cell 2021, 39:1479–1496 e1418.

40. Martin BK, Chin KC, Olsen JC, Skinner CA, Dey A, Ozato K, Ting JP: Induction of MHC class I expression by the MHC class II transactivator CIITA. Immunity 1997, 6:591–600.

41. Grabosch S, Bulatovic M, Zeng F, Ma T, Zhang L, Ross M, Brozick J, Fang Y, Tseng G, Kim E, et al: Cisplatin-induced immune modulation in ovarian cancer mouse models with distinct inflammation profiles. Oncogene 2019, 38:2380–2393.

42. Nguyen EM, Taniguchi H, Chan JM, Zhan YA, Chen X, Qiu J, de Stanchina E, Allaj V, Shah NS, Uddin F, et al: Targeting Lysine-Specific Demethylase 1 Rescues Major Histocompatibility Complex Class I Antigen Presentation and Overcomes Programmed Death-Ligand 1 Blockade Resistance in SCLC. J Thorac Oncol 2022, 17:1014–1031.

43. Hiatt JB, Sandborg H, Garrison SM, Arnold HU, Liao SY, Norton JP, Friesen TJ, Wu F, Sutherland KD, Rienhoff HY, et al: Inhibition of LSD1 with Bomedemstat Sensitizes Small Cell Lung Cancer to Immune Checkpoint Blockade and T-Cell Killing. Clin Cancer Res 2022, 28:4551–4564.

44. Eisenhauer EA, Therasse P, Bogaerts J, Schwartz LH, Sargent D, Ford R, Dancey J, Arbuck S, Gwyther S, Mooney M, et al: New response evaluation criteria in solid tumours: revised RECIST guideline (version 1.1). Eur J Cancer 2009, 45:228–247.

45. Ireland AS, Micinski AM, Kastner DW, Guo B, Wait SJ, Spainhower KB, Conley CC, Chen OS, Guthrie MR, Soltero D, et al: MYC Drives Temporal Evolution of Small Cell Lung Cancer Subtypes by Reprogramming Neuroendocrine Fate. Cancer Cell 2020, 38:60–78 e12.

46. Lepelley A, Della Mina E, Van Nieuwenhove E, Waumans L, Fraitag S, Rice GI, Dhir A, Fremond ML, Rodero MP, Seabra L, et al: Enhanced cGAS-STING-dependent interferon signaling associated with mutations in ATAD3A. J Exp Med 2021, 218.

47. Pietanza MC, Waqar SN, Krug LM, Dowlati A, Hann CL, Chiappori A, Owonikoko TK, Woo KM, Cardnell RJ, Fujimoto J, et al: Randomized, Double-Blind, Phase II Study of Temozolomide in Combination With Either Veliparib or Placebo in Patients With Relapsed-Sensitive or Refractory Small-Cell Lung Cancer. J Clin Oncol 2018, 36:2386–2394.

48. Stanzione M, Zhong J, Wong E, LaSalle TJ, Wise JF, Simoneau A, Myers DT, Phat S, Sade-Feldman M, Lawrence MS, et al: Translesion DNA synthesis mediates acquired resistance to olaparib plus temozolomide in small cell lung cancer. Sci Adv 2022, 8:eabn1229.

49. Lukinovic V, Hausmann S, Roth GS, Oyeniran C, Ahmad T, Tsao N, Brickner JR, Casanova AG, Chuffart F, Benitez AM, et al: SMYD3 Impedes Small Cell Lung Cancer Sensitivity to Alkylation Damage through RNF113A Methylation-Phosphorylation Cross-talk. Cancer Discov 2022, 12:2158–2179.

50. Taniguchi H, Caeser R, Chavan SS, Zhan YA, Chow A, Manoj P, Uddin F, Kitai H, Qu R, Hayatt O, et al: WEE1 inhibition enhances the antitumor immune response to PD-L1 blockade by the concomitant activation of STING and STAT1 pathways in SCLC. Cell Rep 2022, 39:110814.

